# Crowder titrations enable the quantification of driving forces for macromolecular phase separation

**DOI:** 10.1101/2023.07.03.547544

**Authors:** Gaurav Chauhan, Anne Bremer, Furqan Dar, Tanja Mittag, Rohit V. Pappu

## Abstract

Macromolecular solubility is an important contributor to the driving forces for phase separation. Formally, the driving forces in a binary mixture comprising a macromolecule dissolved in a solvent can be quantified in terms of the saturation concentration, which is the threshold macromolecular concentration above which the mixture separates into coexisting dense and dilute phases. Additionally, the second virial coefficient, which measures the effective strength of solvent-mediated intermolecular interactions provides direct assessments of solvent quality. The sign and magnitude of second virial coefficients will be governed by a combination of solution conditions and the nature of the macromolecule of interest. Here, we show, using a combination of theory, simulation, and *in vitro* experiments, that titrations of crowders, providing they are true depletants, can be used to extract the intrinsic driving forces for macromolecular phase separation. This refers to saturation concentrations in the absence of crowders and the second virial coefficients that quantify the magnitude of the incompatibility between macromolecules and the solvent. Our results show how the depletion-mediated attractions afforded by crowders can be leveraged to obtain comparative assessments of macromolecule-specific, intrinsic driving forces for phase separation.

**SIGNIFICANCE:** Phase separation has emerged as a process of significant relevance to sorting macromolecules into distinct compartments, thereby enabling spatial and temporal control over cellular matter. Considerable effort is being invested into uncovering the driving forces that enable the separation of macromolecular solutions into coexisting phases. At its heart, this process is governed by the balance of macromolecule-solvent, inter-macromolecule, and solvent-solvent interactions. We show that the driving forces for phase separation, including the coefficients that measure interaction strengths between macromolecules, can be extracted by titrating the concentrations of crowders that enable macromolecules to phase separate at lower concentrations. Our work paves the way to leverage specific categories of measurements for quantitative characterizations of driving forces for phase separation.

## INTRODUCTION

Macromolecular phase separation has been proposed to be important for enabling spatial and temporal organization of cellular matter through the formation of biomolecular condensates (1-7). Phase separation is governed by the free energy of mixing of a macromolecule or collections of macromolecules in a complex solvent. In a mixture comprising a macromolecule dissolved in a low molecular weight solvent, entropy favors the mixing of macromolecule and solvent degrees of freedom. However, if the interactions of macromolecules and the solvent are energetically incompatible, then the solvent quality is poor (8), and there exists a threshold concentration in a binary mixture, or a solubility product for multicomponent systems (9) beyond which the macromolecular solution separates into two or more coexisting phases (2). For a system comprising two coexisting phases, the dense phase is rich in macromolecules, and the dilute phase is rich in solvent. The simplest mean-field models for phase separation treat the solvent implicitly. Its influence is instead captured through the size disparity between the solvent and macromolecule, and through a parameter χ that quantifies the balance of macromolecule-solvent, inter-macromolecule, and solvent-solvent interactions (10).

Cellular solvents are complex, and they combine the contributions of water molecules, mono-and multivalent ionic species, osmolytes, metabolites, and other macromolecules that act as crowders. Of particular interest is the effect of crowders because they can alter the entropic and energetic contributions to the free energy of mixing. We consider a macromolecular solution comprising finite concentrations of the macromolecule of interest, crowders that are large enough to take up significant space in the solution, ligands, and small molecule components of the complex solvent that include water molecules, ions, and small molecule osmolytes. Such systems are complex *n-*nary mixtures, but for simplicity we shall treat them as pseudo binary systems comprising the macromolecules and the crowders of interest dissolved in implicit, albeit complex solvents. How do crowders affect macromolecular phase separation and why is this important? This is a question of significant interest and relevance. Here, we take on a simpler question pertaining to the effects of synthetic crowders on macromolecular phase separation in pseudo binary mixtures.

Synthetic crowders can be spherical, linear, or branched polymers. They can also be modeled as hard or soft sphere colloidal particles. To zeroth order, crowders are thought of as depletants (11) that interact with one another and with the macromolecules via steric repulsions alone. The non-adsorbing nature of the crowders creates a depletion layer around the macromolecules. As the concentrations of crowders increase, the macromolecules are squeezed together, causing an overlap of their depletion layers. This creates an unbalanced force, in the form of osmotic pressure, and the macromolecules are drawn to one another to offset the unbalanced force. In an ideal regime, the osmotic pressure, being a colligative property, is directly proportional to the concentration of the crowder. Considerable effort and attention have been invested into understanding the effects of crowders on conformational, binding, and phase equilibria of biomacromolecules (12-29). These efforts are motivated in part by the view that cells, which are crowded environments, are also non-ideal milieus, and that thermodynamic non-idealities are likely to have profound effects on conformational, binding, and phase equilibria (30-36).

Our focus is on the use of crowders as enablers of macromolecular phase separation *in vitro*. Molecules presumed to be inert that are used as crowders include dextran, ficoll, and polyethylene glycol (PEG) (37-46). The presumption of inertness is intended to imply that these molecules are likely to be pure depletants that do not adsorb on the surfaces of macromolecules, and that the non-steric interactions, if present, are weak when compared to thermal energy (11, 47-49). The motivations for using synthetic crowders are two-fold: First, it can be challenging to obtain high yields of recombinant proteins and nucleic acids (43, 44). The use of synthetic crowders allows for phase separation to be studied in small volumes and at lower concentrations of macromolecules. Second, crowders are thought to be mimics of the crowded environments in cells, and this assumption has been used as justification for their usage *in vitro* (37, 40, 41, 47, 50).

It is common to read reports of condensate formation in the presence of PEG-8000, the 8 kDa version of PEG. The concentrations of PEG-8000 or other forms of PEG that are used *in vitro* typically range from 1 – 10% weight per volume, although higher concentrations have also been used in specific studies. Recent studies have questioned key assumptions that go into the use of molecules such as PEG-8000 as crowders (51, 52). A key finding of concern is the observation that crowders are not excluded from dense phases. Further, it appears that crowders are not generic depletants that leave the slopes of tie lines unchanged (52). In such a scenario, the crowders become co-scaffolds of phase separation, and the dense phase can be dominated by crowders. The latter is the worst-case scenario if the objective is to study / characterize condensates formed by the macromolecules of interest rather than dense phases that are rich in crowders (52). The situation is exacerbated by the fact that one seldom measures the concentrations of macromolecules or crowders in the dense phase because this can be challenging, although this has now been remedied using new and accessible methods (53). Absent such measurements, the composition of the dense phase and the contributions of crowders to the properties of dense phase remain uncertain.

Our work is motivated by the fact that *in vitro* characterizations of macromolecular phase separation are invaluable for bottom-up reconstitutions of facsimiles of condensates (38, 39, 43, 44, 46, 53-72). Such reconstitutions can help with quantifying the saturation concentration *c*_sat_, defined as the threshold concentration beyond which the system separates into macromolecule-rich and solvent-rich phases that coexist with one another. In a pseudo binary mixture where the solvent is implicit and is designated as component 0, the macromolecule is component 1, and any depletant / crowder is component 2, the parameter of interest is the second virial coefficient, *B*_11_, which quantifies the strength and nature of solvent-mediated interactions between pairs of macromolecules (2). Since *B*_11_ is directly proportional to the Flory χ parameter, it is a measure of the quality of the crowder-free solvent for the macromolecule of interest. Note that phase separation can occur if and only if *B*_11_ is negative (2).

Can one use titrations of crowders to obtain estimates of *B*_11_ and *c*_sat,0_, defined as the intrinsic saturation concentration for a given set of solution conditions in the absence of crowder? The answer is yes, and here we extend the pioneering work of Edmond and Ogston (49, 73) to illustrate how intrinsic driving forces for macromolecular phase separation, i.e., values of *B*_11_ and *c*_sat,0_ can be extracted using titrations of crowder concentrations. Our work combines an adaptation of theoretical work pioneered by Edmond and Ogston (73) with simulations designed to explicitly test the assumptions that go into the theory. Having set up the criteria where the theoretical analysis is applicable, we illustrate its use by deploying it to analyze *in vitro* data for the phase separation of a yeast transcription factor.

## RESULTS

### The Edmond-Ogston model (73) and its usage

The pseudo binary mixture comprises an implicit solvent (component 0), the macromolecules of interest (component 1), and the crowders (component 2). Accounting only for effective two-body interactions, the free energy of mixing in the pseudo binary mixture is written as:

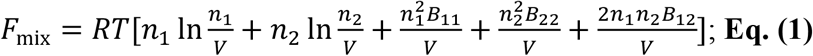

Here, *n*_*i*_ refers to the number of moles of component *i, V* is the volume of composite system, *R* is the ideal gas constant, and *T* is the temperature. The molar concentration of component *i* is given by *c*_*i*_ = *n*_*i*_/*V*. The coefficients *B*_11_ and *B*_22_ are the second virial coefficients, and they have units of association constants. The magnitude and sign of *B*_11_ quantifies the effective strengths and nature of pairwise interactions among macromolecules. Conversely, the magnitude and sign of *B*_12_ quantifies the strength and nature of the effective interactions between the macromolecule and crowder.

The quantities of interest are the chemical potentials μ_1_ and μ_2_, which are the partial molar free energies of the macromolecule and crowder, respectively. For the macromolecule, the chemical potential 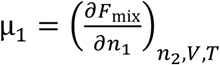 becomes:

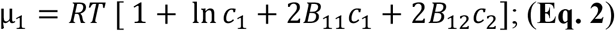

If the crowder plus solvent system is a poor solvent for the macromolecule, then above a threshold concentration of the macromolecule, the system undergoes phase separation. Assuming the crowder is a true depletant, the result of phase separation will be two coexisting phases: a dense phase rich in the macromolecule that coexists with a dilute phase that is rich in the solvent. The macromolecular concentrations in the coexisting phases are set by equalizing the chemical potentials 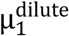 and 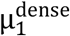 defined as:

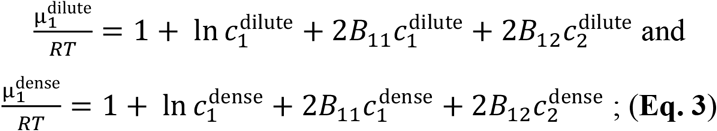

The concentrations of macromolecule and crowder in the coexisting phases are determined by the equalization of the chemical potentials and equalization of the osmotic pressure. Here, we work with chemical potentials of the macromolecule in the dense and dilute phases. Phase equilibrium is established by the requirement that:

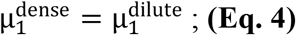

Further, if the crowder is a true depletant, then it should be excluded from the dense phase. It then follows that:

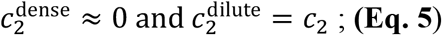

Substituting **Eq. 5** into **Eq. 4**, we obtain the following relationships:

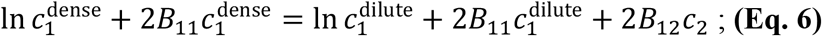

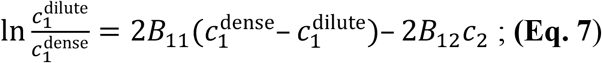

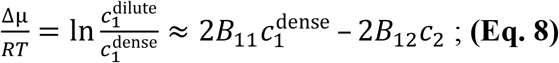

This is essentially an expression for the osmotic pressure, which being a colligative property, is directly proportional to the concentration of the crowder *c*_2_. If the concentrations of the macromolecule in the coexisting dense and dilute phases are known via direct measurement at a series of crowder concentrations, then one can extract *B*_11_ using the fact that *B*_12_ is a slope whose value is constant. Accordingly, we shall consider crowder concentrations designated as *c*_2,*i*_ and *c*_2,*j*_. If we set 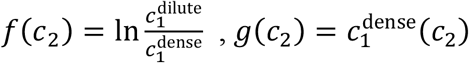, and *a* = 2*B*_11_, then it follows that:

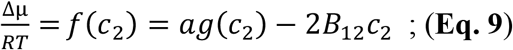

Since *B*_12_ is a constant, it follows that:

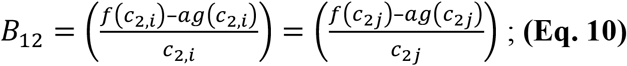

Rearranging the equations, we obtain the following relationships:

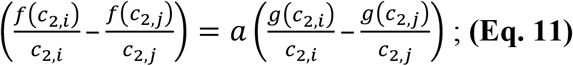

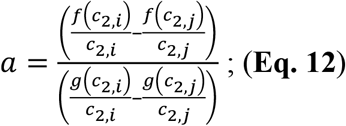

Using pairs of points *i* and *j* along the titration curve that are chosen at random, one can estimate *a* using regression analysis, and this yields an estimate of *B*_11_. Note that this requires knowledge of dense and dilute phase concentrations of the macromolecules as a function of crowder concentration, *c*_2_. It does not require the elaborate measurements that are typically thought to be essential for estimating *B*_11_.

To estimate, the values of *B*_12_ and the concentration of the dilute phase in the absence of crowder, we return to the original formalism of the Edmond-Ogston model. Because 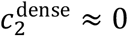, the chemical potential of the macromolecule in the dense phase reduces to:

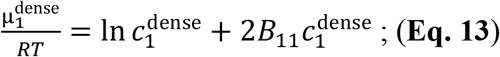

Further, in the limit where the crowder concentration approaches zero, *c*_2_ → 0, we can assume that the macromolecular concentration in the dense phase changes only minimally with the bulk concentration of the crowder *c*_2_. In this limit, we can set 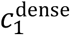 to be a constant, and this implies that 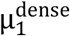 is essentially invariant to changes in crowder concentration. We reiterate that this is valid only for *c*_2_ → 0. Since the macromolecular chemical potentials are equalized at equilibrium, the constancy of the dense phase chemical potential implies that the dilute phase chemical potential will not vary significantly with crowder concentration. Accordingly, for a given value of *c*_2_, in the limit *c*_2_ → 0, if we designate the dilute phase concentration of the macromolecule to be *c*_sat_, and the intrinsic saturation concentration in the absence of crowder as *c*_sat,0_, it follows that:

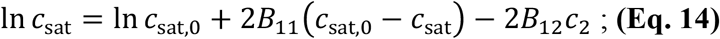

**Eq. 14** is the essence of the Edmond-Ogston model (73). It predicts a log-linear relationship between the macromolecular *c*_sat_ and the crowder concentration *c*_2_. This relationship holds only if the crowder is excluded from the dense phase and the dense phase concentration of the macromolecule remains roughly constant as the crowder concentration changes. Further, and strictly speaking, **Eq. 14** is applicable only if the concentration of the crowder is in the dilute regime where the osmotic pressure scales linearly with *c*_2_. In the semidilute regime, the osmotic pressure scales non-linearly with *c*_2_, and deviations from the log-linear relationship will be governed by the dependence of *B*_12_ on *c*_2_ (74). However, these effects are weakly perturbing as discussed in the Appendix. Therefore, in the limit of *c*_2_ → 0, the log-linear relationship of **Eq. 14** is a reliable expectation for how the macromolecular *c*_sat_ varies with *c*_2_. We used simulations where the macromolecule and crowder are modeled explicitly to show how **Eq. 12** can be used to estimate *B*_11_ and to test the accuracy of **Eq. 14** as an estimator of *c*_sat,0_.

### Simulations for studying the effects of crowding on phase separation

We used coarse-grained, lattice-based simulations to study the role of crowding on macromolecular phase separation and to test the validity of using **Eq. 12** and **Eq. 14**. These simulations use the LaSSI (Lattice Simulation Engine for Stickers and Spacer Interactions) package (75). In the lattice model, the macromolecules are linear sticker and spacer polymers on a lattice (**Fig. 1A**), and the crowders are linear polymer depletants (**Fig. 1B**). The stickers make reversible crosslinks with interaction energies shown in **Fig. 1C**. The explicitly modeled spacers take up room on lattice sites. We examined the effects of crowders by changing their bulk volume fraction ϕ_c_.

**Fig. 1.**
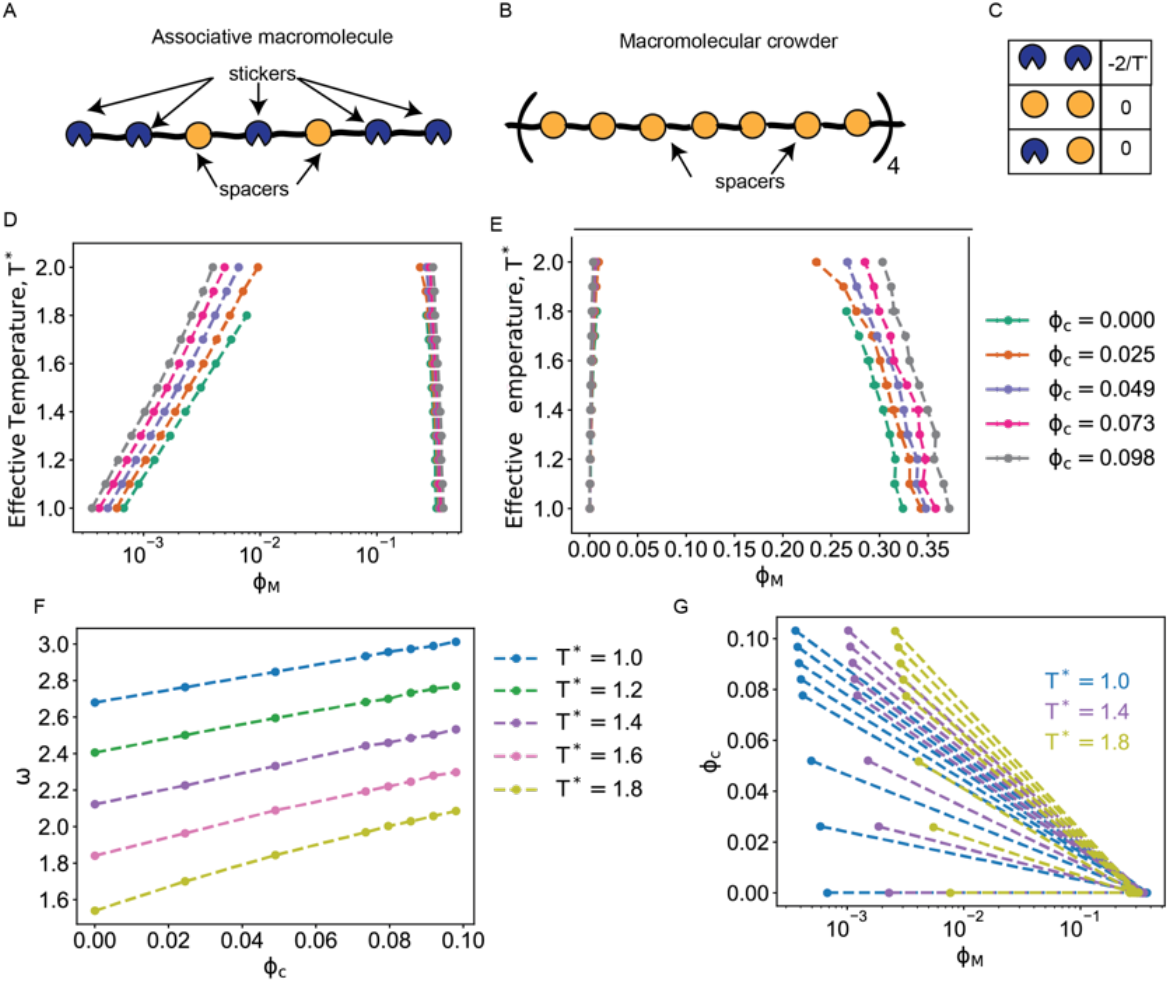
LaSSI simulations were used to probe the effects of crowders on macromolecular phase separation. (A) Schematic of the associative macromolecule with sticker and spacer architecture consisting of five sticker and two spacer beads, with a linker length of two lattice units (l.u.). (B) Schematic of the linear, polymeric crowder comprising 28 beads that gives it a contour length of 28 l.u. (C) Interaction energies between the sticker and spacer beads. The energies are parameterized in units of the reduced simulation temperature, *T*\*. (D) Coexistence curves of the macromolecule computed in the presence of different volume fractions of crowder polymer. (E) Same as D, but with volume fraction along the abscissa shown on a linear scale. (F) Width of the two-phase regime, ω, as a function of ϕ_c_ at different temperatures (shown as different curves). (G) Coexistence curves plotted on a semi-log scale, with crowder volume fractions shown in a linear scale along the ordinate. Results are shown for three different simulation temperatures, *T*\*=1.0, 1.4, 1.8. For a given value of ϕ_c_ and *T*\*, the two-phase regime corresponds to the region between the solid points. The dashed lines are tie lines, and for all non-zero values of ϕ_c_, we note that the slopes of the tie lines are negative, implying that the crowder is excluded from the dense phase. Conversely, the dilute phase volume fraction of the macromolecule decreases with increased crowder concentration, implying depletion of the macromolecule from the dilute phase.

### Increasing ϕ_c_ lowers ϕ_sat_ and increases the width of the two-phase regime

**Figs. 1D** and **1E** show the coexistence curves (binodals) for the macromolecule and its modulation as a function of crowding. The two figures plot the binodals on different scales. Increasing ϕ_c_ from 0 to 0.1 decreases ϕ_sat_ by roughly an order of magnitude across the simulated temperature regime (**Fig. 1D**). Across the range of crowder concentrations, the dense phase concentration ϕ_dense_ increases by a factor of 1.2 (**Fig. 1E**). Note that crowders are unlike ligands that engage in site-specific interactions with macromolecules that undergo phase separation. While ligands dilute the concentrations of macromolecules in the dense phase, even if they interact preferentially with the dense phase (76, 77), crowders, as shown here (**Fig. 1E**), have the opposite effect. Accordingly, an increase in macromolecular concentration within the dense phase is a useful readout of whether component 2 is a crowder or a ligand that binds site-specifically to the macromolecule. Taken together with previous findings (76, 77), our results suggest that endogeneous macromolecules that are not the scaffolds or co-scaffolds of condensates are likely to function as ligands or crowders.

The large, temperature-dependent changes to ϕ_sat_ and the smaller changes to ϕ_dense_ impact the width of the two-phase regime, defined as: ω = ln(ϕ_dense_ / ϕ_sat_). This calculation uses temperature-specific values of ϕ_dense_ and ϕ_sat_ to obtain estimates of ω as a function of the simulation temperature *T*\*. The width of the two-phase regime increases with increase in crowder concentration, and the magnitude of the increase depends on the driving forces for macromolecular phase separation, which decrease with increasing temperature (**Fig. 1F**).

The crowders in the simulation are true depletants that are excluded from the dense phase. This is shown via our analysis of coexistence curves in the plane of macromolecular volume fraction (ϕ_M_) along the abscissa and crowder volume fraction (ϕ_c_) along the ordinate. In these simulations, the contour length of the crowder is more than twice that of the macromolecules (**Fig. 1A** versus **Fig. 1B**). These crowders are clearly excluded from the dense phase as quantified by the fact that the volume fraction of crowder in the dense phase is essentially zero, and the slopes of the tie lines are negative for all non-zero values of ϕ_c_ (**Fig. 1G**).

Next, we analyzed how the dilute phase concentration varies as a function of ϕ_c_ for different temperatures (**Fig. 2A**). We observed a non-linear decrease of ϕ_sat_ as ϕ_c_ increases. Plotting the logarithm of the normalized ϕ_sat_ versus ϕ_c_ reveals signatures of the log-linear relationship between ϕ_sat_ and ϕ_c_ that is predicted by the Edmond-Ogston model (**Fig. 2B**). The volume fraction of the macromolecule in the dense phase increases with the volume fraction of the crowder, and it also decreases with increasing temperature. To unmask how changes to crowder concentration influences the temperature dependence of the volume fraction of the macromolecule in the dense phase, we analyzed the normalized dense phase volume fraction of the macromolecule (**Fig. 2C**). This analysis shows that the macromolecular volume fraction in the dense phase increases, on average across the temperature regime, by a factor of ∼1.2 as the crowder volume fraction increases 0 to 0.1 (**Fig. 2D**). However, the concentration of the macromolecule in the dense phase varies minimally in the limit of small values of ϕ_c_. This suggests that the Edmond-Ogston model of **Eq. 14** might be applicable in this limit.

**Fig. 2.**
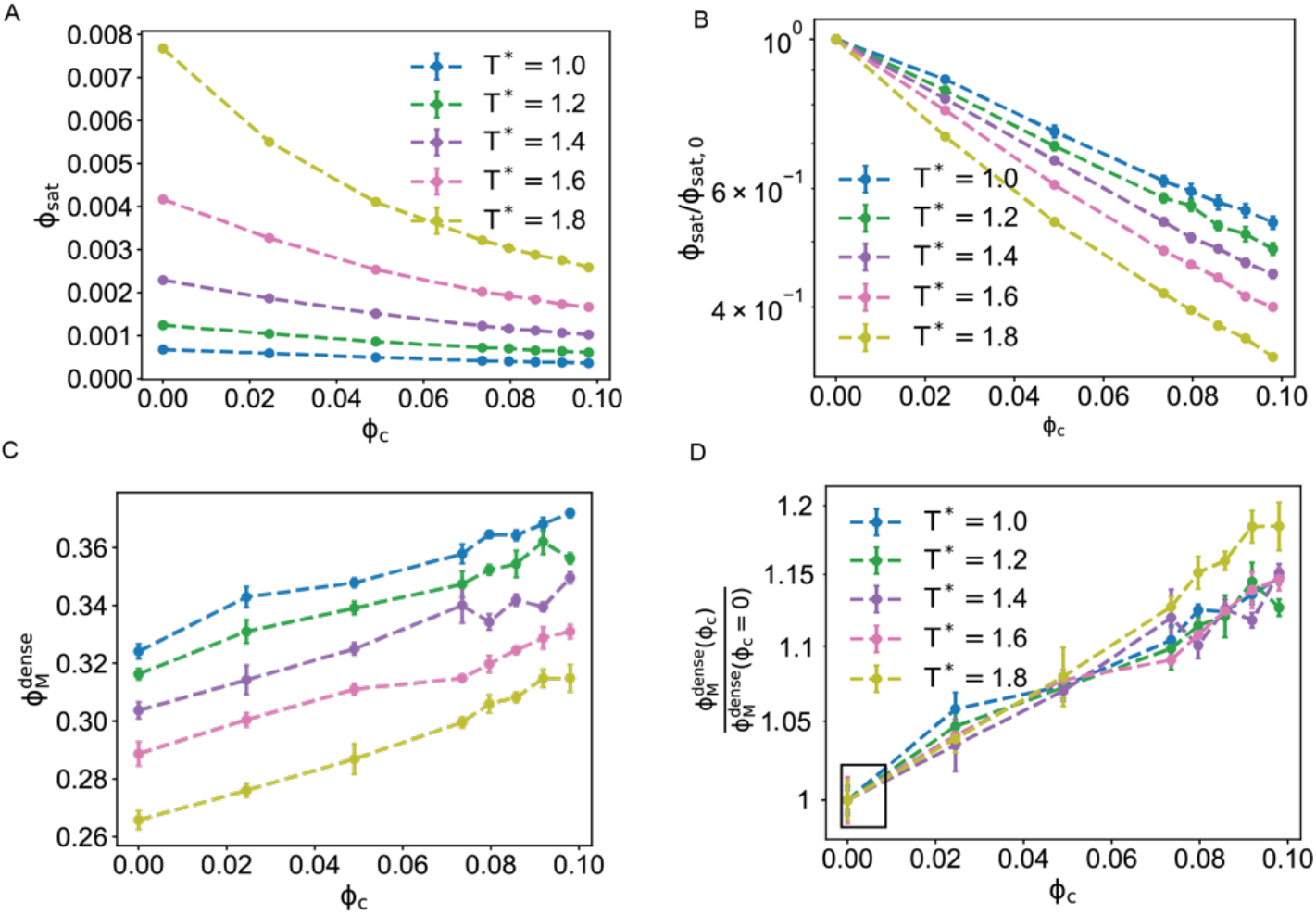
Crowders lower the ϕ_sat_ of macromolecules showing a log-linear relationship between ϕ_sat_ and ϕ_c_. (A) Plot of ϕ_sat_ against ϕ_c_ on a linear scale. (B) Normalized values of ln(ϕ_sat_ / ϕ_sat_|ϕ_c_ =0) plotted against ϕ_c_. Note that the ordinate is on a logarithmic scale, and the abscissa is on a linear scale. (C) The volume fraction of the macromolecule in the dense phase plotted against the volume fraction of the crowder. Each curve corresponds to a different simulation temperature, *T*\*. The colors are identical to those of panel B, which includes the legend for the different temperatures. Note that the driving forces for phase separation weaken with increasing temperature. (D) Normalized volume fraction of the macromolecule in the dense phase plotted against ϕ_c_. The clear box brackets the region where the normalized volume fraction of the macromolecule in the dense phase changes minimally with ϕ_c_.

### Crowding-enabled decreases in saturation concentration depend on crowder-size

Next, we investigated the role of crowder size on macromolecular phase separation. We used the ratio of contour lengths of crowder to macromolecule, defined as (*L*_C_ / *L*_M_), as a measure of the relative sizes of crowders to macromolecules. Irrespective of the value of (*L*_C_ / *L*_M_), the driving forces for phase separation are enhanced in the presence of crowders, as evidenced by the lowering of ϕ_sat_ in the presence of crowders. We quantified how the saturation concentration (**Fig. 3A**) varies with ϕ_c_ for fixed temperature, and different values of *L*_C_ / *L*_M_. Larger crowders are most efficient at lowering the saturation concentration. To understand this result, we computed the radial density profiles of macromolecules and crowders as a function of the radial distance from the center of the dense phase (**Fig. 3B, 3C**). Although the crowder is largely excluded from the dense phase, the degree of exclusion depends on the size of the crowder (**Fig. 3D**). Crowders that are smaller than the macromolecule can partition into the dense phase, whereas crowders that are on a par with or larger than the macromolecules are fully excluded from the dense phase. In a two-phase system, a crowder that is smaller than the macromolecule can passively partition into the dense phase. The driving force for this partitioning derives from the entropy of mixing whereby mixing of the degrees of freedom will contribute to the chemical potentials of the crowder and macromolecule in the dense phase.

**Fig. 3.**
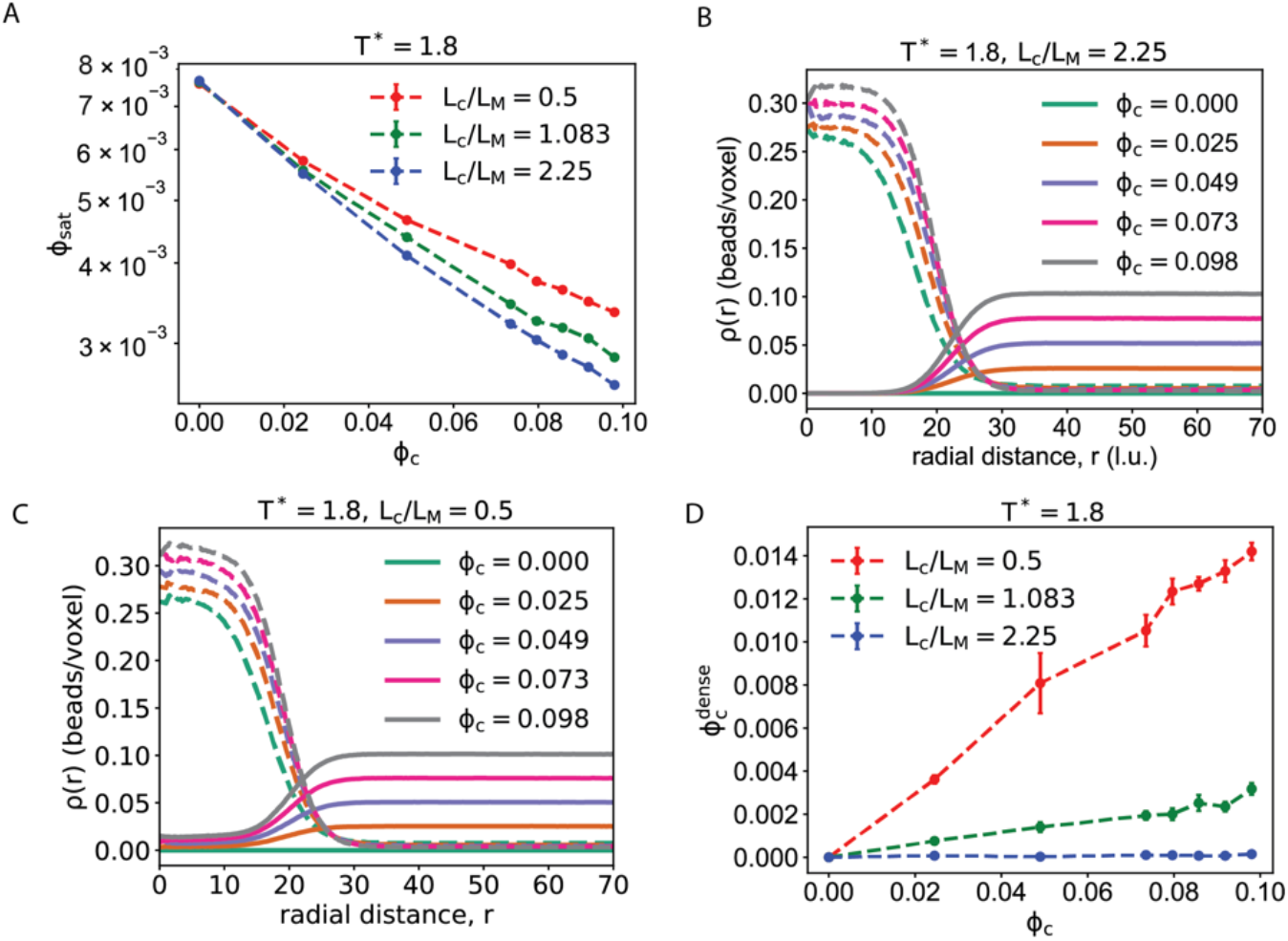
Sizes of crowders affect the driving forces for macromolecular phase separation. (A) Plot of how ϕ_sat_ changes as a function of ϕ_c_ for crowders of different size. The latter is quantified using the ratio (*L*_C_ / *L*_M_) and each curve corresponds to a specific value of this ratio. (B) and (C) Radial density profiles of the macromolecule (dashed lines) and the crowder (solid lines) as a function of the radial distance from the center of mass of the dense phase for a simulation temperature of *T*\* = 1.8 and values of (*L*_C_ / *L*_M_) = 2.25 (B) and 0.5 (C). (D) Volume fraction of the crowder in the dense phase, plotted against the bulk volume fraction of the crowder for a simulation temperature of *T*\* = 1.8 and three different values of (*L*_C_ / *L*_M_).

### *Estimation of B*_*11*_, ϕ_sat,0_, and *B*_12_

Now, we arrive at the central motivation of our work, which is the estimation of parameters that quantify the driving forces for phase separation by leveraging information extracted from titrations of crowders. The protocol we follow involves two distinct steps. The approach prescribed by **Eq. 12** is first applied to estimate *B*_11_. The results are shown as triangles in **Fig. 4A**. The driving forces weaken with increasing temperature, and as a result, *B*_11_ becomes less negative, decreasing in magnitude with increasing *T*\*. Crowders that are on a par with or larger than macromolecules are excluded from the dense phase. For these crowders, the estimates of *B*_11_ are independent of the crowder size. However, the smaller crowder yields values of *B*_11_ that are ∼1.1 times larger in magnitude when we use estimations based on **Eq. 12**. The simplest explanation for the discrepant values that we estimate for *B*_11_ based on small versus large crowders is that the fitting based on Eq. 12 assumes that the dense phase concentrations of crowders are zero, and this is not true for smaller crowders.

**Fig. 4.**
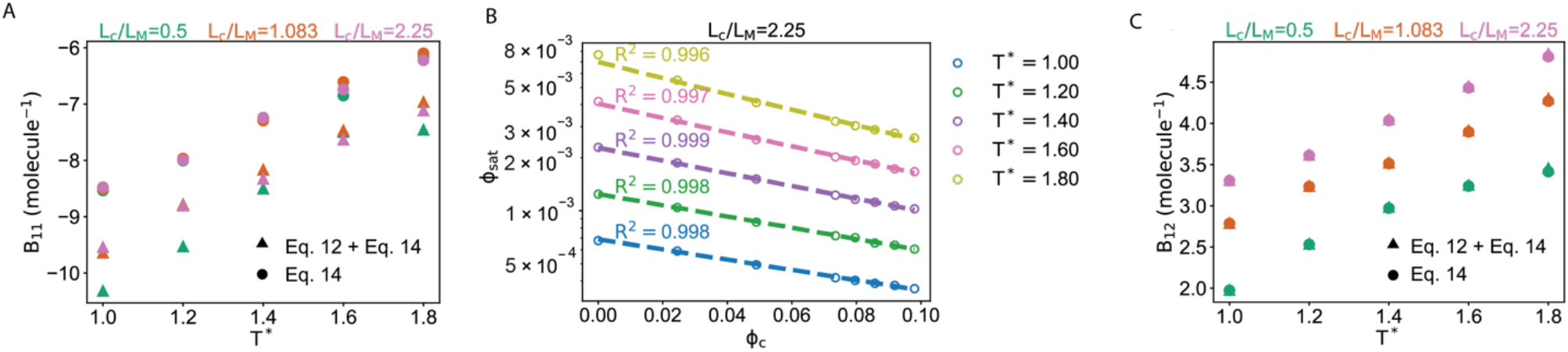
Application of Eq. 12 and Eq. 14 to estimates *B*_11_, *B*_12_, and ϕ_sat,0_. (A) Estimates of *B*_11_ obtained using **Eq. 12** – see circles – and **Eq. 14** – see triangles – for three different crowder sizes. The estimates of *B*_11_ are plotted against simulation temperature *T*\*. The different curves are for different crowders of different sizes. For crowders that are equivalent to or larger in size than the macromolecule, the intrinsic self-association of macromolecules is independent of crowder size and depends only on the simulation temperature. (B) Plots of ln(ϕ_sat_) against ϕ_c_, one curve per temperature (see legend to the right). These plots show the log-linear relationship predicted by the Edmond-Ogston model. Regression analysis, performed using *B*_11_ values extracted using **Eq. 12**, was then deployed to estimate of ϕ_sat,0_. These are shown as intercepts with the ordinate for ϕ_c_ = 0. The values of ϕ_sat,0_ computed directly from the simulations are shown as open circles that correspond to the value of ϕ_sat_ for ϕ_c_ = 0. The dashed lines are fits of ln(ϕ_sat_) to **Eq. 14**. The analysis shows good agreement between the estimated and actual values of ϕ_sat,0._ (C) Plot of estimated values of *B*_12_ against the simulation temperature *T*\*. The different curves are for different crowder sizes.

Next, we fit the simulation results for the variation of ϕ_sat_ as function of ϕ_c_ using **Eq. 14**. We excluded the known values of ϕ_sat,0_ from the fits (**Fig. 4B**). We used the values of *B*_11_ estimated using **Eq. 12** and summarized in **Fig. 4A**. The fits to the Edmond-Ogston log-linear relationship, using known values of *B*_11_, allow us to estimate ϕ_sat,0_ and *B*_12_ (**Fig. 4C**). The values of *B*_12_ increase and become more positive with increasing simulation temperature (**Fig. 4B**). This highlights the stronger impact of crowders on depletion-mediated attractions when the driving forces for macromolecular phase separation are weaker. The drive to increase the concentration of macromolecules in the dilute phase, aided by weaker driving forces for phase separation, will be opposed by the drive to minimize the loss of free volume for the crowders in the dilute phase. The intrinsic self-association of macromolecules, quantified by the macromolecular second virial coefficient *B*_11_, becomes less negative with simulation temperature (**Fig. 4A**).

An overwhelming majority of experiments do not measure the concentrations of macromolecules in dense phases. Accordingly, we asked if the values of *B*_11_, ϕ_sat,0_, and *B*_12_ can be estimated using **Eq. 14** alone. The results (see triangles in **Fig. 4A**) show that the use of **Eq. 14** alone leads to systematic underestimation of the values of *B*_11_, and this is true across the temperature range. The two-body interactions contribute to equalizing the chemical potential of the macromolecule across the dense and dilute phases. This in turn sets the concentrations of the coexisting phases at equilibrium. Our analysis shows that while knowledge of the dilute and dense phase concentrations is required for quantitatively accurate estimation of B_11_ through analysis based on **Eq. 12**, the use of crowder titrations and the simplifications that lead to **Eq. 14** provide at least qualitatively accurate estimates, with an underestimation by a factor of 0.9. This seems like an acceptable amount of error given the simplicity of the Edmond-Ogston model.

### Effects of soft interactions with crowders

Interactions between macromolecules and crowders can be parsed into hard and soft contributions (21, 22). Excluded volume effects come under the rubric of hard contributions. Soft interactions can either be attractive or repulsive in nature. To probe the effects of attractions between crowders and macromolecules, we performed additional LaSSI simulations by incorporating attractive isotropic interactions between crowders and macromolecules. We modulated their strengths to study how crowders with soft, attractive interactions affect the intrinsic and crowding-enabled phase behaviors. For interactions that are weaker than those of the inter-macromolecule interactions, we observed a preservation of the log-linear relationship between ϕ_sat_ and ϕ_c_ (**Fig. 5A**). Previous studies have highlighted the weakening of depletion interactions as the soft, attractive interactions between crowders and macromolecules become stronger (23). Consistent with these studies, we note that the estimated values of *B*_12_ decrease as the attractive interactions between crowders and macromolecules become stronger.

**Fig. 5.**
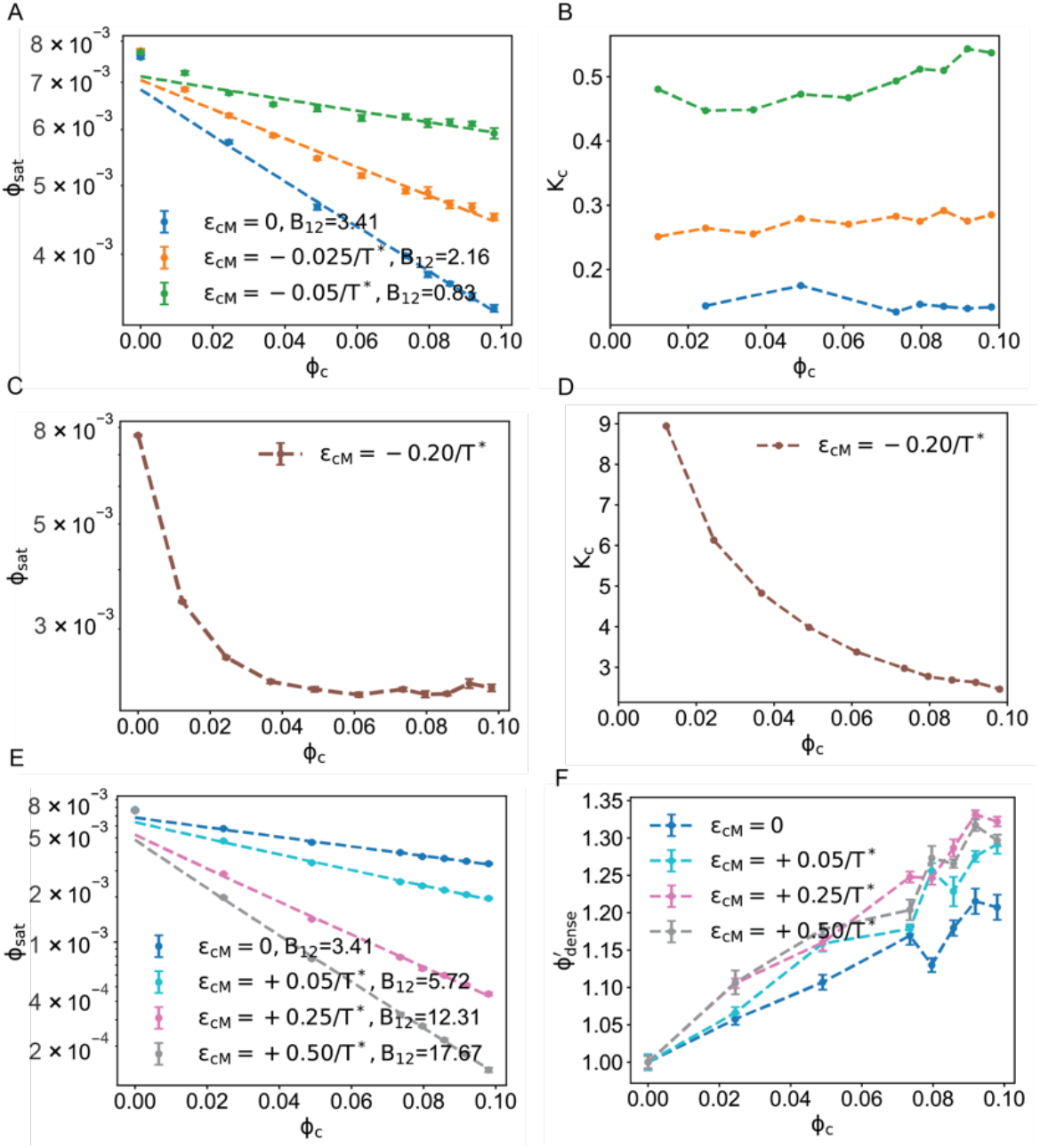
Contributions of soft interactions between crowders and macromolecules. (A) Variation of ϕ_sat_ as a function of ϕ_c_, for different interaction strengths ε_CM_ between the crowder and macromolecule sites. The negative sign denotes attractive interactions. Here, the interaction strengths are weak enough to ensure that the crowders function as depletants. (B) Partition coefficients (*K*_C_) of crowders plotted against ϕ_c_ for different values of ε_CM_. (C) Variation of ϕ_sat_ as a function of ϕ_c_ when the interactions between macromolecule and crowder are strong (ε_CM_ = –0.20/*T*\*). In this scenario, a blend of favorable homotypic interactions among macromolecules and attractive interactions between crowders and macromolecules drive phase separation. (D) Partition coefficient of the crowder into the dense phase plotted as a function of ϕ_c_ for ε_CM_ = –0.20/*T*\*. Note that these values are above unity for all values ϕ_c_ that were investigated. (E) Saturation concentration of the macromolecule as a function of ϕ_c_ for the case of added soft repulsions between crowders and macromolecules. (F) Normalized dense phase volume fraction, ϕ′_dense_, defined as the ratio of the dense phase volume fraction of the macromolecule in the presence versus the absence of crowder as a function of ϕ_c_ for different strengths of soft repulsive interactions between crowders and macromolecules.

Next, we quantified the partition coefficients of the crowders for different values of the interaction strengths between crowders and macromolecules (**Fig. 5B**). The partition coefficient of the crowder, which is the ratio of the concentration of the crowder in the dense versus dilute phase, increases as the crowder-macromolecule interactions increase in strength. However, comparisons between the results in **Fig. 5B** and **Fig. 5A** show that the log-linear relationship between ϕ_sat_ and ϕ_c_ is preserved, even when the soft interactions are attractive, providing the partition coefficient of the crowder is less than unity.

Increasing the strength of the crowder-macromolecule interactions above a threshold value, which is –0.20/*T*\* for the current model, we note that the log-linear relationship between ϕ_sat_ and ϕ_c_ is fundamentally altered (**Fig. 5C**). The partition coefficient of the crowder is now greater than one, and there is a non-linear relationship between the partition coefficient and ϕ_c_ (**Fig. 5D**). In this scenario, a blend of homotypic interactions between the macromolecule and heterotypic interactions between the crowder and macromolecule jointly drive phase separation. The crowder now functions as a co-scaffold instead of being a depletant.

Next, we investigated the effects of soft repulsions that add to the excluded volume effects of crowders. Such soft repulsions can arise if solvent molecules adsorb on the surfaces of crowders. These super-depletant crowders lower the saturation concentrations, more so than the purely excluded volume crowders (**Fig. 5E**). The inclusion of soft repulsions between crowders and macromolecules enhances depletion-mediated attractions because the crowders require larger free volumes within the dilute phase. As noted for the purely excluded volume crowders, the concentration of the macromolecule in the dense phase increases by a factor of 1.1 – 1.3, depending on the bulk concentration of the crowder. The dense phase concentrations of macromolecules change with ϕ_c_ but are essentially insensitive to the strengths of the soft repulsions between crowders and macromolecules (**Fig. 5F**).

### *Errors in estimates of* ϕ_sat,0_

We quantified the errors in estimates of ϕ_sat,0_ obtained using Eq. 14 with *B*_11_ values obtained using **Eq. 12** (**Fig. 6**). The error increases as the ratio of *B*_12_-to-|*B*_11_| increases. Note that *B*_12_ increases and the magnitude of B_11_ decreases with increasing temperature. Therefore, the ratio of *B*_12_-to-|*B*_11_| increases with increasing temperature. Our results show that the error in estimates of ϕ_sat,0_ will increase as the intrinsic driving forces for phase separation are weakened. As a reminder, these estimates are obtained using a combination of **Eq. 12** and **Eq. 14** that use information from measurements of how the concentrations of macromolecules in dilute and dense phases change with crowder titrations. Recent studies have shown that the value of ϕ_sat,0_ will increase by 2-3 orders of magnitude over the range of experimentally accessible temperatures (55). When compared to this range of variation, errors of 8-10% that define the estimates of ϕ_sat,0_ from titrations of crowders can be viewed as being reasonably reliable.

**Fig. 6:**
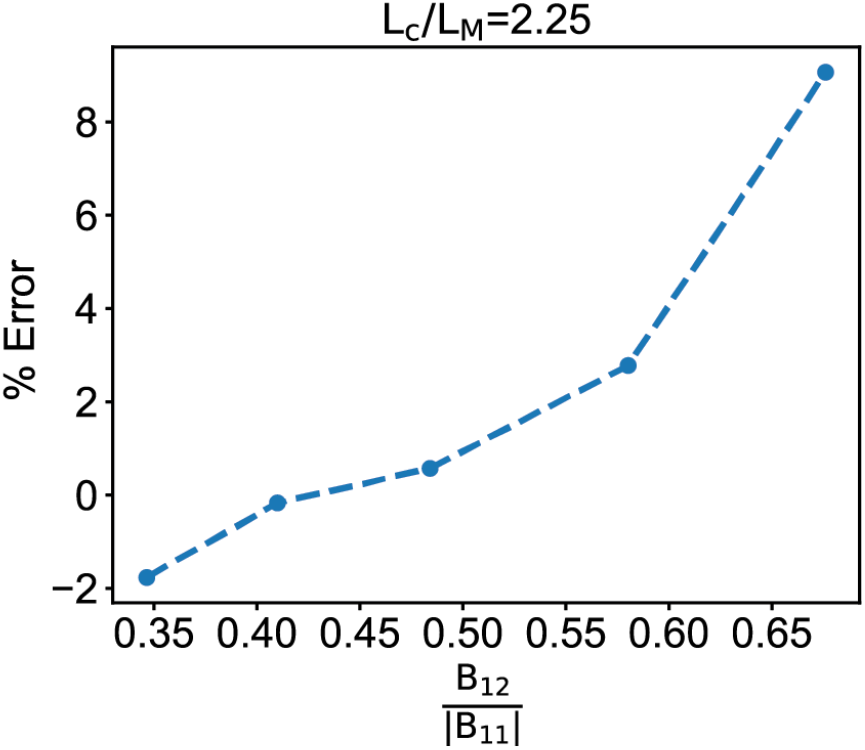
Errors in estimates of ϕ_sat,0_ using the combination of Eq. 12 and E1q. 13 increase with increases to the ratio of *B*_12_-to-|*B*_11_|. This analysis was performed for results from simulations that use crowders with an (*L*_C_ / *L*_M_) value of 2.25. The % error was computed as 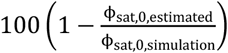.

### Summary of findings from theory and simulation

Overall, our simulations lead to the following predictions for the effects of crowders on macromolecular phase separation. If the crowders interact purely via excluded volume effects, then they will lower ϕ_sat_ through depletion-mediated interactions. An ideal crowder is one that is excluded from the dense phase. The inclusion of soft attractions between the macromolecule and crowders will preserve the log-linear relationship between the crowder concentration dependent value of ϕ_sat_ and the crowder volume fraction ϕ_c_ providing the partition coefficient of the crowder is less than unity. Soft repulsions will enhance the depletion effect. Analysis of how the concentrations of macromolecules in the dense and dilute phases change with crowder titrations can be used to extract values of *B*_11_, ϕ_sat,0_, and *B*_12_. We demonstrate this using *in vitro* experiments as described next.

### Testing predictions using in vitro experiments

PEG-8000 is commonly used as a crowder to induce macromolecular phase separation *in vitro*. We illustrate how titrations of PEG-8000 can be used to extract the intrinsic driving forces for a short form of the yeast transcription factor Gcn4. The sequence of the protein we studied (see Methods) spans the central, intrinsically disordered activation domain (residues 101-141) from *S. cerevisiae* connected by a short (GS)_4_-linker to the DNA-binding domain of Gcn4 (residues 222-281). In a 20 mM HEPES buffer, at pH 7.3 with 150 mM potassium acetate and 2 mM DTT the intrinsic saturation concentration of wild-type (WT) Gcn4 is in the millimolar range. We used a recently introduced HPLC assay (53) to measure the dilute and dense phase concentrations of WT Gcn4 as a function of the concentration of PEG-8000.

WT Gcn4 forms micron-scale condensates in the presence of PEG-8000 (**Fig. 7A**). We performed titrations of crowder concentrations from 2.5 w/v % to 15 w/v %. The concentration of WT Gcn4 in the dense phase changes minimally with crowder concentration (**Fig. 7B**). The measured values of *c*_sat_ decrease with increasing concentrations of crowder (**Fig. 7C**). The data shown in **Figs. 7B** and **7C** were used to extract *B*_11_ using **Eq. 12**, and the estimate of *B*_11_ was used in regression analysis based on **Eq. 14** to obtain estimates of *c*_sat,0_ and *B*_12_. The residuals we obtain from the fitting are shown in **Fig. 7D**. The estimates of *c*_sat,0_ obtained using the two-step process that combines analysis based on **Eq. 12** and **Eq. 14** are compared to those obtained using **Eq. 14** alone (**Fig. 7C**). In general, we obtained consistent estimates for c_sat,0_ using both methods. However, the estimates for *B*_11_ are off by a factor of two, suggesting, in line with the simulation results, that measurements of dense phase concentrations of macromolecules are helpful because they provide additional constraints.

**Fig. 7.**
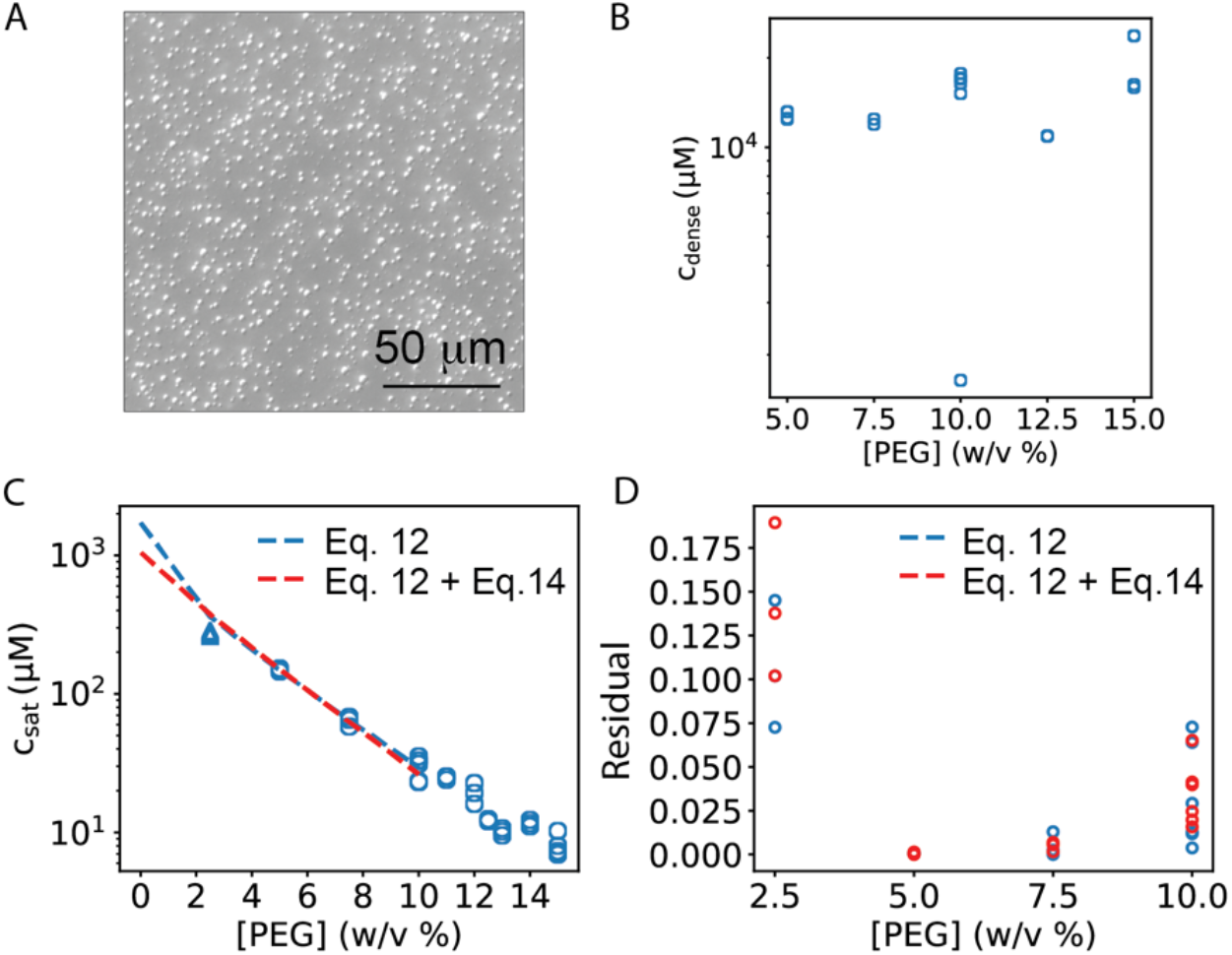
Illustration of how crowder titrations *in vitro* can be used for extracting information regarding intrinsic driving forces for macromolecular phase separation. (A) WT Gcn4 forms micron-scale condensates *in vitro* in the presence of PEG-8000. The image shown here is from differential interference contrast (DIC) microscopy for WT Gcn4 in the presence of 10 w/v % of PEG-8000. (B) The concentration of WT Gcn4 in the dense phase ranges from a minimum of ∼11 mM to a maximum of ∼15 mM, although the average value is ∼12 mM across a range of crowder concentrations. (C) Crowder titrations were used in conjunction with **Eq. 12** and **Eq. 14** and based on **Eq. 12** alone, to estimate *c*_sat,0_. Using the combination of **Eq. 12** and **Eq. 14**, we obtained values of *B*_11_ = –0.13 /mM, *B*_12_ = 0.14 / mM, and *c*_sat,0_ = 1.05 mM. Using **Eq. 14** alone, we obtained values of *B*_11_ = –0.29 /mM, *B*_12_ = 0.12 / mM, and *c*_sat,0_ = 1.72 mM. These estimates are reasonably similar across the two methods, and the results highlight how crowder titrations can be used to glean insights regarding the intrinsic driving forces for macromolecular phase separation. (D) For each concentration of the crowder, PEG-800, the residuals of the fits shown in panel (C) are plotted against [PEG]. These were computed as: 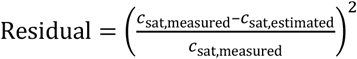.

Our simulation results suggest that crowders, if they are large enough, should be excluded from dense phases. We tested these predictions using PEG molecules of two different molecular weights. We used a colorimetric assay to quantify the concentration of crowders in the coexisting dilute phase versus the bulk concentration of crowders. The latter is the input concentration of the crowder. The ratio of concentration of PEG-8000 in the dilute phase versus the bulk concentration is shown in **Fig. 8A**. A value of unity would indicate that the concentration of PEG in the dilute phase is equal to that of the input bulk concentration. We observed that the concentration of PEG-8000 is larger in the dilute phase than that of the input bulk concentration. This suggests that the crowder is depleted from the dense phase. We also quantified the ratio of crowder concentration in the dilute phase to bulk concentration of crowder for lower molecular weight crowder namely, PEG-2000. This concentration ratio is less than one, implying that there is accumulation of PEG-2000 in the dense phase. It also suggests that there are favorable interactions between the smaller crowder and the macromolecules in the dense phase.

**Fig. 8:**
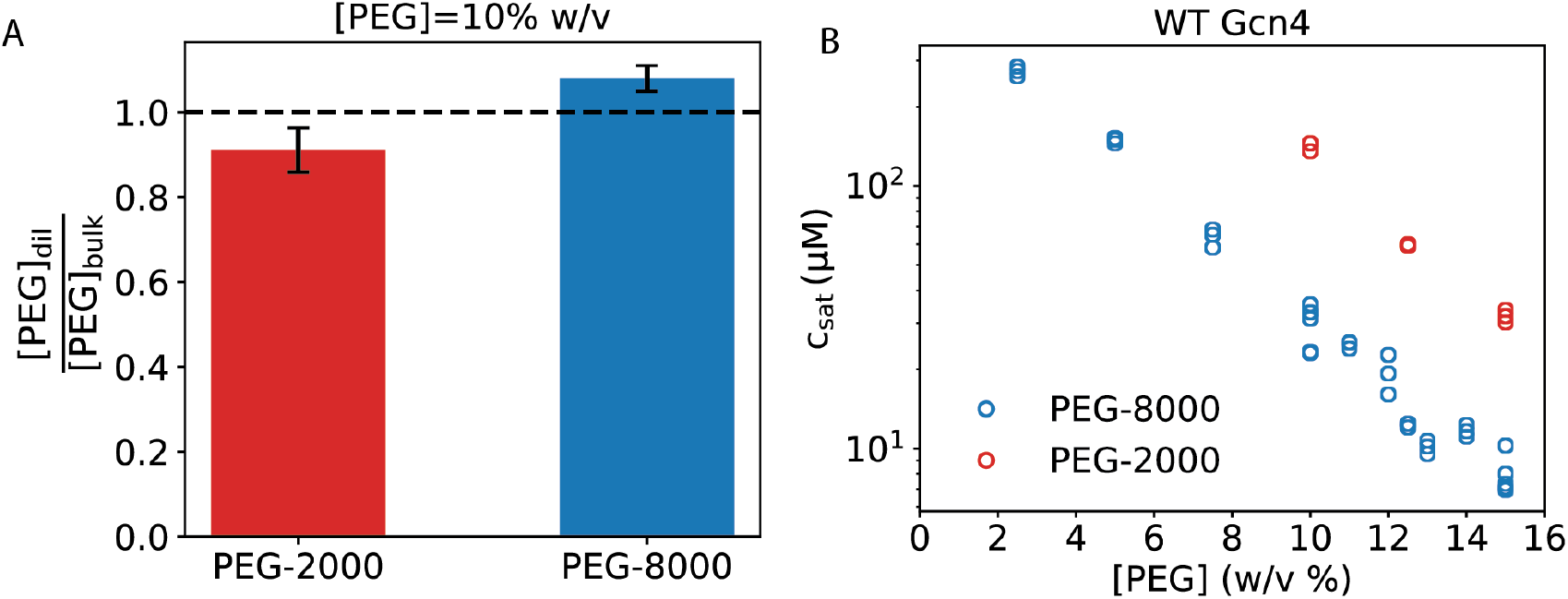
Assessments of the impact of two different crowders on phase separation of WT Gcn4. (A) Ratio of PEG concentration in the dilute phase to the input PEG concentration for PEG-2000 and PEG-8000. The measurements were performed at a crowder weight to volume ratio of 10%. (B) Measured saturation concentrations for WT Gcn4 as a function of PEG concentration for PEG-2000 and PEG-8000.

Next, we compared how the saturation concentrations of WT Gcn4 evolve as a function of crowder concentrations for PEG-2000 and PEG-8000. For equivalent weight to volume ratios of the crowders, the measured *c*_sat_ values are higher for the smaller versus larger crowder. Despite the apparent appearance of the log-linear relationship between *c*_sat_ and crowder concentration, we cannot use the relationship in **Eq. 14** to extract the intrinsic *c*_sat_ because the colorimetric assay clearly shows partitioning of PEG-2000 into the dense phase formed by Gcn4. These results highlight the joint importance of titrating crowder concentrations *and* ensuring that the crowders are excluded from the dense phase.

The recent studies of Stringer et al., (78) showed that low molecular weight PEG molecules have weak attractions to macromolecules, a feature that is lost as the molecular weights increase. The weak attractions are attributable to the more prominent contributions of end groups of low molecular weight PEG molecules. These end group effects are overshadowed by the excluded volume effects of higher molecular weight PEG molecules. The findings from our measurements affirm predictions from simulations and highlight the importance of choosing crowders that are equivalent to or larger than the macromolecules whose phase behaviors are being investigated.

## DISCUSSION

We show how the Edmond-Ogston model (73) leads to a framework for quantitative and comparative analysis of the driving forces for macromolecular phase separation. The framework is captured in the structure of **Eq. 12** and **Eq. 14**. Simulations based on a lattice model were used to test the overall validity of and assumptions underlying the Edmond-Ogston model (73). While the assumptions are not perfect, they are robust, and the model was found to be useful for describing observations from simulations. Based on these results, we deployed the Edmond-Ogston model to analyze results from *in vitro* measurements of phase separation for Gcn4. These systems require high protein concentrations for phase separation to be realized *in vitro* thereby serving as archetypes of systems that are induced to undergo phase separation by the presence of crowders.

Using titrations of PEG-8000, we were able to extract the intrinsic values of *c*_sat_, designated as *c*_sat,0_, and the second virial coefficients. These experiments illustrate how theory can be deployed to analyze experimental data. While the original Edmond-Ogston model based on **Eq. 14** does not require the measurement of macromolecular concentrations in the dense phase, the combined use of **Eq. 12** and **Eq. 14** requires measurements of concentrations in the dense phase. The simulations and *in vitro* experiments show similar discrepancies, typically a factor of two, between estimates of parameters that we obtain using either **Eq. 12** and **Eq. 14** or **Eq. 14** alone. The overall inference is that crowder titrations can be used to extract intrinsic driving forces for phase separation providing the crowders are true depletants and the partition coefficients into the dense phase are below unity. Our results suggest that values of *B*_11_, which are key parameters that are seldom measured, are accessible using readily deployable crowder titrations (45).

What are the criteria that need to be satisfied for the Edmond-Ogston model to be used? First, the crowder must be chosen so its size is on a par with or larger than that of the macromolecule of interest. This implies that one must not use PEG-8000 as a one size fits all generic crowder. Instead, larger PEG molecules become essential when studying proteins of higher molecular weights. Second, suitable assays such as the colorimetric measurements used in this work or inferences regarding the slopes of tie lines used in recent studies must be deployed to ensure that the crowder is largely excluded from the dense phase. Alternatively, one could perform measurements of the partition coefficient of the crowder to ensure that these coefficients remain below one across the range of crowder concentrations used. Third, systematic titrations of the concentration of the crowder must be performed, and the macromolecular *c*_sat_ must be measured at the different concentrations of crowders.

Synthetic crowders are routinely used *in vitro* to drive macromolecular phase separation and enable the formation of stable, micron-sized macrophases. These macrophases, irrespective of whether they are purely viscous liquids, viscoelastic materials, or solids, will grow via coarsening (79) driven by coalescence (80), Ostwald ripening (81), or some combination of the two processes. Often one uses characterizations *in vitro*, that also include crowders, for comparative assessments of driving forces for phase separation. For example, one might quantify the amounts of two types of proteins needed to drive phase separation in the presence of a fixed amount of PEG-8000. These data are then used to stipulate that one protein is better at driving phase separation when compared to the other. Such conclusions are fraught with concerns because the effects of crowders might be different for the two proteins. Our studies, which combine theory, simulations and experiments help establish key “boundary conditions” for the use of crowders.

Here, we investigated the effects of crowders on a system that separates into two coexisting phases. However, most macromolecules can form multiple coexisting phases that are defined by distinct crystalline or liquid-crystalline order parameters (82). Further, certain systems likely form stable microphases such as micelles or emulsions (83, 84). A single *c*_sat_ will not define the crossing of multiple stable or metastable phase boundaries. Crowders will likely act as selective agents that tilt the thermodynamic balance toward a macrophase, microphase, or mesophase that ensures global stability for the macromolecule, plus crowder, plus solvent system. These considerations will mandate the need for separate investigations to probe the formation of microphases or phases defined by different structural symmetries for the macromolecules of interest. A case in point is that of alpha synuclein, which was recently shown to undergo macrophase separation in the presence of PEG-8000 and at higher concentrations in the absence of PEG-8000 (85). However, a recent computational study showed that alpha synuclein, in the absence of crowders, undergoes microphase separation and that the microphases can form a system-spanning percolated network (86). In this case, the crowding-enabled macrophase separation at lower protein concentrations might be an artefact of depletion-mediated attractions that override the intrinsic drive for microphase separation.

The effects of crowders on the wide range of micro- and mesophases, and pre-percolation clusters (87) remains unknown at this juncture. This is worth bearing in mind since microphases and pre-percolation cluster may become destabilized by synthetic crowders. Whether this is the case *in cellulo* will require new and different types of investigations. A simple test would be to assess whether microphases can form, and if they do, then their sizes will stay mostly fixed as protein concentration increases, although the overall abundance will increase. This contrasts with macrophase separation, which will invariably feature some form of coarsening or at least heterogeneity of condensate sizes.

## METHODS

### Computational methods

All simulations were carried out using the lattice simulation engine LaSSI (75). Each simulation consisted of 2000 macromolecules, which were divided into two sets such that the sticker-sticker interactions are only between complementary pairs of molecules. All other interactions of the macromolecule are of the excluded volume variety. The architecture chosen for the macromolecular system is reminiscent of the poly-SH3 + poly-PRM and / or poly-SUM + poly-SIM systems that have been studied by Rosen and coworkers (88, 89). The layout of the lattice model, with a combination of explicit and implicit spacers, which determine whether the excluded volume is that of the lattice or zero, is adapted from the work of Harmon et al., (90, 91). The crowders are modeled as linear polymers, akin to the PEG system (92, 93). In the simulations, the crowders are either pure depletants, modeled as self-avoiding polymers, or their excluded volume contributions are augmented with isotropic, non-site-specific soft interactions that can be attractive or repulsive. Attractive interactions will lower the depletion layer because it enables some degree of adsorption of the crowder onto the macromolecule. Repulsive interactions enhance the depletion effect by adding to the excluded volume effect. The number of crowder polymers were varied to change the volume fraction occupied by crowders. The box side for the cubic simulation box was L=100 lattice units.

The simulation procedure was similar to that used by Ruff et al. (76). Briefly, all the molecules are randomly placed on the lattice without overlap. All the anisotropic interactions between molecules are turned off and a biasing potential is applied to each bead which is meant to push molecules towards the center of the simulation box. The system is then exponentially cooled down to *T*\*=1 from *T*\*=100 for 5 × 10^6^ Monte Carlo steps. This leads to the formation of one large condensate if the conditions correspond to the two-phase regime. Then the system is run for each cycle for 2.5 × 10^9^ MC steps and the temperature is discontinuously changed by Δ*T* =0.10 to correspond to temperature for each cycle. Three independent sets of simulation are performed for each solution conditions. The following relative frequency of the MC moves is used:

| Move type | Relative frequency |
| --- | --- |
| Local | 5000 |
| Bond Rotation | 500 |
| Co-local | 1000 |
| Multi-local | 500 |
| Snake | 500 |
| Chain Translation | 1000 |
| Chain Pivot | 1000 |
| Double Pivot | 1000 |
| Limited Cluster | 30 |
| Large Cluster | 3 |

The coexisting dense and dilute phase volume fractions are obtained by constructing the radial density profile from the center of mass of the system as is shown before (75, 94). Firstly, the number histograms *H(r*_*i*_) are generated from the center-of-mass of the system with a bin-width of 0.25 and are normalized by the number histograms of lattice sites (*H*_*0*_*(r*_*i*_*i)* for a cubic box of dimensions L=100 with periodic boundary conditions and same bin width of 0.25. The density profile is thus *ρ(r*_*i*_) = *H(r*_*i*_)/*H*_*0*_*(r*_*i*_). Concentrations of the macromolecules and crowders in dense phases were computed by averaging over the first ten bins of the radial density profiles whereas dilute phase concentrations were computed by averaging over the last hundred bins.

### *Regression analysis based on the joint use of* **Eq. 12** *and* **Eq. 14**

We used dense and dilute phase volume fractions at each crowder volume fraction to perform fits to **Eq. 12** to obtain *B*_11_. Dense and dilute phase volume fractions for a pair of crowder volume fractions were drawn at random. This method of sampling with replacement was repeated for a total of 5× 10^4^ times. We used the distribution of estimates of *B*_11_ to obtain the mean value for each pair of crowder volume fractions considered. Next, we used the non-linear least squares method in `scipy` (https://scipy.org) to fit dilute phase volume fraction and crowder volume fraction to **Eq. 14** to obtain values for *B*_12_ and 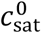. The non-linear least squares method was used exclusively when the fits were based on **Eq. 14** alone.

## Experimental methods

### Details of protein constructs

We used a short version of the canonical yeast transcription factor Gcn4 spanning the central activation domain (residues 101-141) from *S. cerevisiae* (UniProt: P03069) connected by a short (GS)_4_-linker to the DNA-binding domain of Gcn4 (residues 222–281). The sequences were synthesized as previously described (55) including an N-terminal TEV cleavage site followed by the gene of interest. The proteins were expressed in BL21pLysS (DE3) *E. coli* cells and purified from inclusion bodies. The strategy for protein purification was adapted from the published work of Bremer et al., (55). Since Gcn4 contains a cysteine residue, 5 mM 2-Mercaptoethanol was added to all buffers during the purification process. The sequence of WT Gcn4 is shown below.

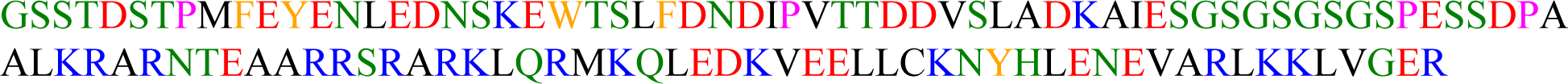

The purified proteins were stored in 6 M GdmHCl (pH 7.3), 20 mM HEPES, 1 mM TCEP at 4°C until the buffer exchange into the phase separation buffer.

### Phase separation assay

The buffer exchange into the phase separation buffer (20 mM HEPES pH 7.3, 2 mM DTT) was achieved by using 2 mL Zeba Spin Desalting columns with 7 MWCO (ThermoFisher). Phase separation was induced by adding PEG and potassium acetate so that the final buffer was 20 mM HEPES (pH 7.3), 150 mM potassium acetate, 0-15% PEG, 2 mM DTT. The samples were incubated at 20°C for 20 min, then centrifuged for 5 min at 12,000 rpm to separate the dilute and dense phase. A defined volume of the dilute phase was then directly transferred to an HPLC vial. We used a positive displacement pipette to transfer 2 μL of the dense phase and diluted it 1000-fold with 6 M GdmHCl (pH 7.3), 20 mM HEPES, 2 mM DTT prior to measurements by analytical HPLC.

### Quantification of saturation and dense phase concentration using analytical HPLC

The protein samples were run on a Waters HPLC system with an Autosampler (Waters 2707), a Binary HPLC Pump (Waters 1525) and a dual-channel UV/Visible Detector (Waters 2489) to determine the saturation (*c*_sat_) and dense (*c*_dense_) phase concentration for Gcn4. We used a C18 reverse-phase column for all samples. The samples were eluted from the column by running an acetonitrile gradient (Alfa Aesar) in H_2_O + 0.1% TFA (Sigma-Aldrich). To determine the saturation and dense phase concentration of Gcn4 we measured a standard curve. The standard curve was fitted to equation 1 from the work of Bremer et al., (53). At least three replicates per condition were measured.

### Colorimetric assay to determine PEG concentration

We used a colorimetric assay developed by Nag et al., (95) and applied it to quantify the partitioning of PEG in the dilute phase. Specifically, 0.5 mL of ammonium ferrothiocyanate and 0.5 mL of chloroform were mixed in a 1.5 mL microfuge tube and 50 μL of the light phase was added to it. The samples were mixed vigorously for 30 min, and the tubes were centrifuged at 3000 g for 2 min. The upper phase was removed and the absorbance of the lower chloroform layer containing PEG was recorded at 510 nm using a cuvette with a 10 mm pathlength. A standard curve of known PEG 8K and PEG 2K were recorded in parallel. At least three replicates per condition were measured.

## APPENDIX Considerations to account for when the crowder concentration is in the semidilute regime

The theoretical approach that we adapted and deployed is applicable in the regime where the osmotic pressure is idea, and the concentration of the crowder is in the dilute regime. In the semidilute regime, the osmotic pressure varies as 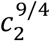 (74). Further, in experiments it is not uncommon for crowder titrations to span the dilute and semidilute regimes. Crowders, especially if they are flexible polymers such as PEG, will overlap with one another in semidilute regime (96). The size of a polymer (measured in terms of the radius of gyration *R*_*g*_) scales as 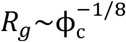 as a function of the volume fraction ϕ_c_ of the crowder. Accordingly, *B*_12_ in **Eq. 1** must be replaced by *B*_12_(ϕ_c_) in the semidilute regime, since *B*_12_ is not constant but will instead be a function of relative volume fraction 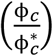 of the crowder where *ϕ*^*\**^ is the overlap volume fraction. For a flexible polymer like crowder molecule with only volume exclusion, the second virial coefficient scales as *B*_22_ *∼*(*R*_*g*,2_)^3^ (97). Similarly, the coefficient *B*_12_ scales as *B ∼*(*R*_*g*,1_ + *R*_*g*,2_)^3^ (73, 98). The radius of gyration below the overlap concentration *ϕ*^*\**^ for the crowder is constant (*R*_*g*,2_(0)) and the coefficient *B*_12_ can be simplified as: 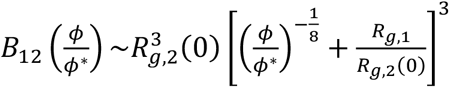. In the limit of *R*_*g*,1_>>*R*_*g*,2_, *B*_12_ is constant in the semidilute regime. In the opposite limit of 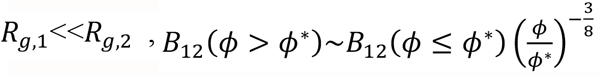 and in the regime of 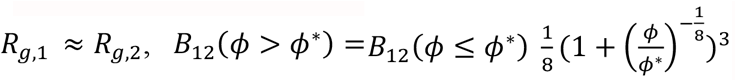. Hence, it follows overlap the regime, *B*_12_ decreases weakly with increased crowder concentration above the. Accordingly, the intricacies of the semidilute regime are unlikely to have a material impact on the analysis introduced in this work, especially if one uses a combination of **Eq. 12** and **Eq. 14** for independent assessments of *B*_11_, *c*_sat,0_, and *B*_12_.

## AUTHOR CONTRIBUTIONS

Conceptualization, G.C., and R.V.P. Design and implementation of the analysis, G.C., and R.V. P. Design of the simulations, G.C., and F.D. Implementation of the simulations and analysis of the simulation results, G.C. Design, and implementation of the *in vitro* measurements, A.B., and T.M. Analysis of the in vitro data, G.C., and A.B. Preparation of the figures, G.C., A.B., and R.V.P. Writing of the manuscript, G.C., and R.V.P. Editing and revision of the manuscript, G.C., A.B., T.M., and R.V.P.

## ACKNOWLEGMENTS

This work was supported by grants from the US National Institutes of Health (R01NS121114 to R.V.P and T.M) and the St. Jude Research Collaborative on the Biology and Biophysics of RNP granules (R.V.P. and T.M). We thank Kresten Lindorff-Larsen for helpful discussions.

## REFERENCES

1. Banani, S. F., H. O. Lee, A. A. Hyman, and M. K. Rosen. 2017. Biomolecular condensates: organizers of cellular biochemistry. Nature reviews. Molecular cell biology 18:285–298.

2. Pappu, R. V., S. R. Cohen, F. Dar, M. Farag, and M. Kar. 2023. Phase Transitions of Associative Biomacromolecules. Chemical Reviews 10.1021/acs.chemrev.2c00814.

3. Alberti, S., and A. A. Hyman. 2021. Biomolecular condensates at the nexus of cellular stress, protein aggregation disease and ageing. Nature Reviews Molecular Cell Biology 22:196–213.

4. Shin, Y., and C. P. Brangwynne. 2017. Liquid phase condensation in cell physiology and disease. Science 357:eaaf4382.

5. Mathieu, C., R. V. Pappu, and J. P. Taylor. 2020. Beyond aggregation: Pathological phase transitions in neurodegenerative disease. Science 370:56–60.

6. Choi, J.-M., A. S. Holehouse, and R. V. Pappu. 2020. Physical Principles Underlying the Complex Biology of Intracellular Phase Transitions. Annual Review of Biophysics 49:107–133.

7. Dignon, G. L., W. Zheng, Y. C. Kim, R. B. Best, and J. Mittal. 2018. Sequence determinants of protein phase behavior from a coarse-grained model. PLoS Computational Biology 14:e1005941.

8. Pappu, R. V., X. Wang, A. Vitalis, and S. L. Crick. 2008. A polymer physics perspective on driving forces and mechanisms for protein aggregation. Archives of Biochemistry and Biophysics 469:132–141.

9. Chattaraj, A., M. L. Blinov, and L. M. Loew. 2021. The solubility product extends the buffering concept to heterotypic biomolecular condensates. eLife 10:e67176.

10. Flory, P. J. 1942. Thermodynamics of High Polymer Solutions. The Journal of Chemical Physics 10:51–61.

11. Asakura, S., and F. Oosawa. 1958. Interaction between particles suspended in solutions of macromolecules. Journal of Polymer Science 33:183–192.

12. Akabayov, B., S. R. Akabayov, S.-J. Lee, G. Wagner, and C. C. Richardson. 2013. Impact of macromolecular crowding on DNA replication. Nature Communications 4:1615.

13. Sharp, K. A. 2015. Analysis of the size dependence of macromolecular crowding shows that smaller is better. Proceedings of the National Academy of Sciences 112:7990–7995.

14. Anderson, C. F., and J. M. Thomas Record. 1995. Salt-Nucleic Acid Interactions. Annual Review of Physical Chemistry 46:657–700.

15. Cayley, S., and M. T. Record Jr. 2004. Large changes in cytoplasmic biopolymer concentration with osmolality indicate that macromolecular crowding may regulate protein–DNA interactions and growth rate in osmotically stressed Escherichia coli K-12. Journal of Molecular Recognition 17:488–496.

16. Konopka, M. C., J. C. Weisshaar, and M. T. Record. 2007. Chapter Twenty-Seven - Methods of Changing Biopolymer Volume Fraction and Cytoplasmic Solute Concentrations for In Vivo Biophysical Studies. In Methods in Enzymology. D. Häussinger, and H. Sies, editors. Academic Press. 487–504.

17. Soranno, A., I. Koenig, M. B. Borgia, H. Hofmann, F. Zosel, D. Nettels, and B. Schuler. 2014. Single-molecule spectroscopy reveals polymer effects of disordered proteins in crowded environments. Proceedings of the National Academy of Sciences 111:4874–4879.

18. Cubuk, J., and A. Soranno. 2022. Macromolecular Crowding and Intrinsically Disordered Proteins: A Polymer Physics Perspective. ChemSystemsChem 4:e202100051.

19. König, I., A. Soranno, D. Nettels, and B. Schuler. 2021. Impact of In-Cell and In-Vitro Crowding on the Conformations and Dynamics of an Intrinsically Disordered Protein. Angewandte Chemie International Edition 60:10724–10729.

20. Zosel, F., A. Soranno, K. J. Buholzer, D. Nettels, and B. Schuler. 2020. Depletion interactions modulate the binding between disordered proteins in crowded environments. Proceedings of the National Academy of Sciences 117:13480–13489.

21. Miklos, A. C., C. Li, N. G. Sharaf, and G. J. Pielak. 2010. Volume Exclusion and Soft Interaction Effects on Protein Stability under Crowded Conditions. Biochemistry 49:6984–6991.

22. Sarkar, M., C. Li, and G. J. Pielak. 2013. Soft interactions and crowding. Biophysical Reviews 5:187–194.

23. Speer, S. L., C. J. Stewart, L. Sapir, D. Harries, and G. J. Pielak. 2022. Macromolecular Crowding Is More than Hard-Core Repulsions. Annual Review of Biophysics 51:267–300.

24. Miller, Cayla M., Young C. Kim, and J. Mittal. 2016. Protein Composition Determines the Effect of Crowding on the Properties of Disordered Proteins. Biophysical Journal 111:28–37.

25. Kim, Y. C., and J. Mittal. 2013. Crowding Induced Entropy-Enthalpy Compensation in Protein Association Equilibria. Physical Review Letters 110:208102.

26. Kang, H., P. A. Pincus, C. Hyeon, and D. Thirumalai. 2015. Effects of Macromolecular Crowding on the Collapse of Biopolymers. Physical Review Letters 114:068303.

27. Cheung, M. S., D. Klimov, and D. Thirumalai. 2005. Molecular crowding enhances native state stability and refolding rates of globular proteins. Proceedings of the National Academy of Sciences 102:4753–4758.

28. Pincus, D. L., and D. Thirumalai. 2009. Crowding Effects on the Mechanical Stability and Unfolding Pathways of Ubiquitin. The Journal of Physical Chemistry B 113:359–368.

29. Denesyuk, N. A., and D. Thirumalai. 2011. Crowding Promotes the Switch from Hairpin to Pseudoknot Conformation in Human Telomerase RNA. Journal of the American Chemical Society 133:11858–11861.

30. Minton, A. P. 2005. Influence of macromolecular crowding upon the stability and state of association of proteins: Predictions and observations. Journal of Pharmaceutical Sciences 94:1668–1675.

31. Ellis, R. J. 2001. Macromolecular crowding: obvious but underappreciated. Trends in Biochemical Sciences 26:597–604.

32. Mourão Márcio A., Joe B. Hakim, and S. Schnell. 2014. Connecting the Dots: The Effects of Macromolecular Crowding on Cell Physiology. Biophysical Journal 107:2761–2766.

33. Rivas, G., and A. P. Minton. 2016. Macromolecular Crowding In Vitro, In Vivo, and In Between. Trends in Biochemical Sciences 41:970–981.

34. Delarue, M., G. P. Brittingham, S. Pfeffer, I. V. Surovtsev, S. Pinglay, K. J. Kennedy, M. Schaffer, J. I. Gutierrez, D. Sang, G. Poterewicz, J. K. Chung, J. M. Plitzko, J. T. Groves, C. Jacobs-Wagner, B. D. Engel, and L. J. Holt. 2018. mTORC1 Controls Phase Separation and the Biophysical Properties of the Cytoplasm by Tuning Crowding. Cell 174:338-349.e320.

35. Sang, D., T. Shu, C. F. Pantoja, A. Ibáñez de Opakua, M. Zweckstetter, and L. J. Holt. 2022. Condensed-phase signaling can expand kinase specificity and respond to macromolecular crowding. Molecular Cell 82:3693-3711.e3610.

36. Jalihal, A. P., S. Pitchiaya, L. Xiao, P. Bawa, X. Jiang, K. Bedi, A. Parolia, M. Cieslik, M. Ljungman, A. M. Chinnaiyan, and N. G. Walter. 2020. Multivalent Proteins Rapidly and Reversibly Phase-Separate upon Osmotic Cell Volume Change. Molecular Cell 79:978-990.e975.

37. Woodruff, J. B., B. Ferreira Gomes, P. O. Widlund, J. Mahamid, A. Honigmann, and A. A. Hyman. 2017. The Centrosome Is a Selective Condensate that Nucleates Microtubules by Concentrating Tubulin. Cell 169:1066-1077.e1010.

38. Guillen-Boixet, J., A. Kopach, A. S. Holehouse, S. Wittmann, M. Jahnel, R. Schlussler, K. Kim, I. Trussina, J. Wang, D. Mateju, I. Poser, S. Maharana, M. Ruer-Gruss, D. Richter, X. Zhang, Y. T. Chang, J. Guck, A. Honigmann, J. Mahamid, A. A. Hyman, R. V. Pappu, S. Alberti, and T. M. Franzmann. 2020. RNA-Induced Conformational Switching and Clustering of G3BP Drive Stress Granule Assembly by Condensation. Cell 181:346–361 e317.

39. Wang, J., J.-M. Choi, A. S. Holehouse, H. O. Lee, X. Zhang, M. Jahnel, S. Maharana, R. Lemaitre, A. Pozniakovsky, D. Drechsel, I. Poser, R. V. Pappu, S. Alberti, and A. A. Hyman. 2018. A Molecular Grammar Governing the Driving Forces for Phase Separation of Prion-like RNA Binding Proteins. Cell 174:688-699.e616.

40. Mitrea, D. M., J. A. Cika, C. B. Stanley, A. Nourse, P. L. Onuchic, P. R. Banerjee, A. H. Phillips, C.-G. Park, A. A. Deniz, and R. W. Kriwacki. 2018. Self-interaction of NPM1 modulates multiple mechanisms of liquid–liquid phase separation. Nature Communications 9:842.

41. Marianelli, A. M., B. M. Miller, and C. D. Keating. 2018. Impact of macromolecular crowding on RNA/spermine complex coacervation and oligonucleotide compartmentalization. Soft Matter 14:368–378.

42. Petronilho, E. C., M. M. Pedrote, M. A. Marques, Y. M. Passos, M. F. Mota, B. Jakobus, G. d. S. d. Sousa, F. Pereira da Costa, A. L. Felix, G. D. S. Ferretti, F. P. Almeida, Y. Cordeiro, T. C. R. G. Vieira, G. A. P. de Oliveira, and J. L. Silva. 2021. Phase separation of p53 precedes aggregation and is affected by oncogenic mutations and ligands. Chemical Science 12:7334–7349.

43. Li, W., J. Hu, B. Shi, F. Palomba, M. A. Digman, E. Gratton, and H. Jiang. 2020. Biophysical properties of AKAP95 protein condensates regulate splicing and tumorigenesis. Nature cell biology 22:960–972.

44. Shi, B., W. Li, Y. Song, Z. Wang, R. Ju, A. Ulman, J. Hu, F. Palomba, Y. Zhao, J. P. Le, W. Jarrard, D. Dimoff, M. A. Digman, E. Gratton, C. Zang, and H. Jiang. 2021. UTX condensation underlies its tumour-suppressive activity. Nature 597:726–731.

45. Annunziata, O., N. Asherie, A. Lomakin, J. Pande, O. Ogun, and G. B. Benedek. 2002. Effect of polyethylene glycol on the liquid–liquid phase transition in aqueous protein solutions. Proceedings of the National Academy of Sciences 99:14165–14170.

46. Molliex, A., J. Temirov, J. Lee, M. Coughlin, A. P. Kanagaraj, H. J. Kim, T. Mittag, and J. P. Taylor. 2015. Phase separation by low complexity domains promotes stress granule assembly and drives pathological fibrillization. Cell 163:123–133.

47. Marenduzzo, D., K. Finan, and P. R. Cook. 2006. The depletion attraction: an underappreciated force driving cellular organization. Journal of Cell Biology 175:681–686.

48. Hansen, J., P. N. Pusey, P. B. Warren, A. G. Yodh, K. Lin, J. C. Crocker, A. D. Dinsmore, R. Verma, and P. D. Kaplan. 2001. Entropically driven self–assembly and interaction in suspension. Philosophical Transactions of the Royal Society of London. Series A: Mathematical, Physical and Engineering Sciences 359:921–937.

49. Ogston, A. G. 1970. On the interaction of solute molecules with porous networks. The Journal of Physical Chemistry 74:668–669.

50. Ferrolino, M. C., D. M. Mitrea, J. R. Michael, and R. W. Kriwacki. 2018. Compositional adaptability in NPM1-SURF6 scaffolding networks enabled by dynamic switching of phase separation mechanisms. Nature Communications 9:5064.

51. André, A. A. M., N. A. Yewdall, and E. Spruijt. 2023. Crowding-induced phase separation and gelling by co-condensation of PEG in NPM1-rRNA condensates. Biophysical Journal 122:397–407.

52. Qian, D., T. J. Welsh, N. A. Erkamp, S. Qamar, J. Nixon-Abell, G. Krainer, P. S. George-Hyslop, T. C. T. Michaels, and T. P. J. Knowles. 2022. Tie-lines reveal interactions driving heteromolecular condensate formation. Physical Review X 12:041038.

53. Bremer, A., A. E. Posey, M. B. Borgia, W. M. Borcherds, M. Farag, R. V. Pappu, and T. Mittag. 2022. Quantifying Coexistence Concentrations in Multi-Component Phase-Separating Systems Using Analytical HPLC. Biomolecules 12:1480.

54. Martin, E. W., A. S. Holehouse, I. Peran, M. Farag, J. J. Incicco, A. Bremer, C. R. Grace, M. Soranno, R. V. Pappu, and T. Mittag. 2020. Valence and patterning of aromatic residues determine the phase behavior of prion-like domains. Science 367:694–699.

55. Bremer, A., M. Farag, W. M. Borcherds, I. Peran, E. W. Martin, R. V. Pappu, and T. Mittag. 2022. Deciphering how naturally occurring sequence features impact the phase behaviours of disordered prion-like domains. Nature Chemistry 14:196–207.

56. Jawerth, L., E. Fischer-Friedrich, S. Saha, J. Wang, T. Franzmann, X. Zhang, J. Sachweh, M. Ruer, M. Ijavi, S. Saha, J. Mahamid, A. A. Hyman, and F. Julicher. 2020. Protein condensates as aging Maxwell fluids. Science 370:1317–1323.

57. Maharana, S., J. Wang, D. K. Papadopoulos, D. Richter, A. Pozniakovsky, I. Poser, M. Bickle, S. Rizk, J. Guillén-Boixet, T. M. Franzmann, M. Jahnel, L. Marrone, Y.-T. Chang, J. Sterneckert, P. Tomancak, A. A. Hyman, and S. Alberti. 2018. RNA buffers the phase separation behavior of prion-like RNA binding proteins. Science 360:918–921.

58. Kroschwald, S., M. C. Munder, S. Maharana, T. M. Franzmann, D. Richter, M. Ruer, A. A. Hyman, and S. Alberti. 2018. Different Material States of Pub1 Condensates Define Distinct Modes of Stress Adaptation and Recovery. Cell Reports 23:3327–3339.

59. Franzmann, T. M., M. Jahnel, A. Pozniakovsky, J. Mahamid, A. S. Holehouse, E. Nuske, D. Richter, W. Baumeister, S. W. Grill, R. V. Pappu, A. A. Hyman, and S. Alberti. 2018. Phase separation of a yeast prion protein promotes cellular fitness. Science 359.

60. Patel, A., H. O. Lee, L. Jawerth, S. Maharana, M. Jahnel, M. Y. Hein, S. Stoynov, J. Mahamid, S. Saha, T. M. Franzmann, A. Pozniakovski, I. Poser, N. Maghelli, L. A. Royer, M. Weigert, E. W. Myers, S. Grill, D. Drechsel, A. A. Hyman, and S. Alberti. 2015. A liquid-to-solid phase transition of the ALS protein FUS accelerated by disease mutation. Cell 162:1066–1077.

61. Mateju, D., T. M. Franzmann, A. Patel, A. Kopach, E. E. Boczek, S. Maharana, H. O. Lee, S. Carra, A. A. Hyman, and S. Alberti. 2017. An aberrant phase transition of stress granules triggered by misfolded protein and prevented by chaperone function. The EMBO journal:e201695957.

62. Yang, P., C. Mathieu, R. M. Kolaitis, P. Zhang, J. Messing, U. Yurtsever, Z. Yang, J. Wu, Y. Li, Q. Pan, J. Yu, E. W. Martin, T. Mittag, H. J. Kim, and J. P. Taylor. 2020. G3BP1 Is a Tunable Switch that Triggers Phase Separation to Assemble Stress Granules. Cell 181:325–345 e328.

63. Taylor, N. O., M. T. Wei, H. A. Stone, and C. P. Brangwynne. 2019. Quantifying Dynamics in Phase-Separated Condensates Using Fluorescence Recovery after Photobleaching. Biophysical Journal 117:1285–1300.

64. Freibaum, B. D., J. Messing, P. Yang, H. J. Kim, and J. P. Taylor. 2021. High-fidelity reconstitution of stress granules and nucleoli in mammalian cellular lysate. Journal of Cell Biology 220.

65. Zhang, H., S. Elbaum-Garfinkle, E. M. Langdon, N. Taylor, P. Occhipinti, A. A. Bridges, C. P. Brangwynne, and A. S. Gladfelter. 2015. RNA controls PolyQ protein phase transitions. Molecular Cell 60:220–230.

66. Boeynaems, S., E. Bogaert, D. Kovacs, A. Konijnenberg, E. Timmerman, A. Volkov, M. Guharoy, M. De Decker, T. Jaspers, V. H. Ryan, A. M. Janke, P. Baatsen, T. Vercruysse, R.-M. Kolaitis, D. Daelemans, J. P. Taylor, N. Kedersha, P. Anderson, F. Impens, F. Sobott, J. Schymkowitz, F. Rousseau, N. L. Fawzi, W. Robberecht, P. Van Damme, P. Tompa, and L. Van Den Bosch. 2017. Phase Separation of C9orf72 Dipeptide Repeats Perturbs Stress Granule Dynamics. Molecular Cell 65:1044-1055.e1045.

67. Boija, A., I. A. Klein, B. R. Sabari, A. Dall’Agnese, E. L. Coffey, A. V. Zamudio, C. H. Li, K. Shrinivas, J. C. Manteiga, N. M. Hannett, B. J. Abraham, L. K. Afeyan, Y. E. Guo, J. K. Rimel, C. B. Fant, J. Schuijers, T. I. Lee, D. J. Taatjes, and R. A. Young. 2018. Transcription Factors Activate Genes through the Phase-Separation Capacity of Their Activation Domains. Cell 175:1842–1855 e1816.

68. Guo, Y. E., J. C. Manteiga, J. E. Henninger, B. R. Sabari, A. Dall’Agnese, N. M. Hannett, J. H. Spille, L. K. Afeyan, A. V. Zamudio, K. Shrinivas, B. J. Abraham, A. Boija, T. M. Decker, J. K. Rimel, C. B. Fant, T. I. Lee, Cisse, II, P. A. Sharp, D. J. Taatjes, and R. A. Young. 2019. Pol II phosphorylation regulates a switch between transcriptional and splicing condensates. Nature 572:543–548.

69. Sabari, B. R., A. Dall’Agnese, A. Boija, I. A. Klein, E. L. Coffey, K. Shrinivas, B. J. Abraham, N. M. Hannett, A. V. Zamudio, J. C. Manteiga, C. H. Li, Y. E. Guo, D. S. Day, J. Schuijers, E. Vasile, S. Malik, D. Hnisz, T. I. Lee, I. I. Cisse, R. G. Roeder, P. A. Sharp, A. K. Chakraborty, and R. A. Young. 2018. Coactivator condensation at super-enhancers links phase separation and gene control. Science 361:eaar3958.

70. Henninger, J. E., O. Oksuz, K. Shrinivas, I. Sagi, G. LeRoy, M. M. Zheng, J. O. Andrews, A. V. Zamudio, C. Lazaris, N. M. Hannett, T. I. Lee, P. A. Sharp, I. I. Cissé, A. K. Chakraborty, and R. A. Young. 2021. RNA-Mediated Feedback Control of Transcriptional Condensates. Cell 184:207-225.e224.

71. Schuster, B. S., G. L. Dignon, W. S. Tang, F. M. Kelley, A. K. Ranganath, C. N. Jahnke, A. G. Simpkins, R. M. Regy, D. A. Hammer, M. C. Good, and J. Mittal. 2020. Identifying sequence perturbations to an intrinsically disordered protein that determine its phase-separation behavior. Proceedings of the National Academy of Sciences 117:11421–11431.

72. Kelley, F. M., B. Favetta, R. M. Regy, J. Mittal, and B. S. Schuster. 2021. Amphiphilic proteins coassemble into multiphasic condensates and act as biomolecular surfactants. Proceedings of the National Academy of Sciences 118:e2109967118.

73. Edmond, E., and A. G. Ogston. 1968. An approach to the study of phase separation in ternary aqueous systems. Biochemical Journal 109:569–576.

74. Cohen, J. A., R. Podgornik, P. L. Hansen, and V. A. Parsegian. 2009. A Phenomenological One-Parameter Equation of State for Osmotic Pressures of PEG and Other Neutral Flexible Polymers in Good Solvents. The Journal of Physical Chemistry B 113:3709–3714.

75. Choi, J.-M., F. Dar, and R. V. Pappu. 2019. LASSI: A lattice model for simulating phase transitions of multivalent proteins. PLoS computational biology 15:e1007028.

76. Ruff, K. M., F. Dar, and R. V. Pappu. 2021. Ligand effects on phase separation of multivalent macromolecules. Proceedings of the National Academy of Sciences USA 118:e2017184118.

77. Ruff, K. M., F. Dar, and R. V. Pappu. 2021. Polyphasic linkage and the impact of ligand binding on the regulation of biomolecular condensates. Biophysics Reviews 2:021302.

78. Stringer, M. A., J. Cubuk, J. J. Incicco, D. Roy, K. B. Hall, M. D. Stuchell-Brereton, and A. Soranno. 2023. Excluded Volume and Weak Interactions in Crowded Solutions Modulate Conformations and RNA Binding of an Intrinsically Disordered Tail. The Journal of Physical Chemistry B https://doi.org/10.1021/acs.jpcb.3c02356.

79. Bray, A. J. 1994. Theory of phase-ordering kinetics. Advances in Physics 43:357–459.

80. Berry, J., S. C. Weber, N. Vaidya, M. Haataja, and C. P. Brangwynne. 2015. RNA transcription modulates phase transition-driven nuclear body assembly. Proceedings of the National Academy of Sciences 112:E5237–5245.

81. Lifshitz, I. M., and V. V. Slyozov. 1961. The kinetics of precipitation from supersaturated solid solutions. Journal of Physics and Chemistry of Solids 19:35–50.

82. Azzari, P., M. Bagnani, and R. Mezzenga. 2021. Liquid–liquid crystalline phase separation in biological filamentous colloids: nucleation, growth and order–order transitions of cholesteric tactoids. Soft Matter 17:6627–6636.

83. Yamazaki, T., T. Yamamoto, H. Yoshino, S. Souquere, S. Nakagawa, G. Pierron, and T. Hirose. 2021. Paraspeckles are constructed as block copolymer micelles. The EMBO journal 40:e107270.

84. Ruff, K. M., T. S. Harmon, and R. V. Pappu. 2015. CAMELOT: A machine learning approach for coarse-grained simulations of aggregation of block-copolymeric protein sequences. The Journal of Chemical Physics 143:243123.

85. Ray, S., N. Singh, R. Kumar, K. Patel, S. Pandey, D. Datta, J. Mahato, R. Panigrahi, A. Navalkar, S. Mehra, L. Gadhe, D. Chatterjee, A. S. Sawner, S. Maiti, S. Bhatia, J. A. Gerez, A. Chowdhury, A. Kumar, R. Padinhateeri, R. Riek, G. Krishnamoorthy, and S. K. Maji. 2020. α-Synuclein aggregation nucleates through liquid–liquid phase separation. Nature Chemistry 12:705–716.

86. Das, S., and M. Muthukumar. 2022. Microstructural Organization in α-Synuclein Solutions. Macromolecules 55:4228–4236.

87. Kar, M., F. Dar, T. J. Welsh, L. Vogel, R. Kuhnemuth, A. Majumdar, G. Krainer, T. M. Franzmann, S. Alberti, C. M. Seidel, A. A. Hyman, and R. V. Pappu. 2022. Phase separating RNA binding proteins form heterogeneous distributions of clusters in subsaturated solutions. Proceedings of the National Academy of Sciences 119:e2202222119.

88. Li, P., S. Banjade, H. C. Cheng, S. Kim, B. Chen, L. Guo, M. Llaguno, J. V. Hollingsworth, D. S. King, S. F. Banani, P. S. Russo, Q. X. Jiang, B. T. Nixon, and M. K. Rosen. 2012. Phase transitions in the assembly of multivalent signalling proteins. Nature 483:336–340.

89. Banani, S. F., A. M. Rice, W. B. Peeples, Y. Lin, S. Jain, R. Parker, and M. K. Rosen. 2016. Compositional Control of Phase-Separated Cellular Bodies. Cell 166:651–663.

90. Harmon, T. S., A. S. Holehouse, M. K. Rosen, and R. V. Pappu. 2017. Intrinsically disordered linkers determine the interplay between phase separation and gelation in multivalent proteins. eLife 6:30294.

91. Harmon, T. S., A. S. Holehouse, and R. V. Pappu. 2018. Differential solvation of intrinsically disordered linkers drives the formation of spatially organized droplets in ternary systems of linear multivalent proteins. New Journal of Physics 20:045002.

92. Choi, E., J. Mondal, and A. Yethiraj. 2014. Coarse-Grained Models for Aqueous Polyethylene Glycol Solutions. The Journal of Physical Chemistry B 118:323–329.

93. Trosel, Y., L. P. Gregory, V. K. Booth, and A. Yethiraj. 2023. Diffusion NMR and Rheology of a Model Polymer in Bacterial Cell Lysate Crowders. Biomacromolecules 24:2469–2478.

94. Farag, M., S. R. Cohen, W. M. Borcherds, A. Bremer, T. Mittag, and R. V. Pappu. 2022. Condensates of disordered proteins have small-world network structures and interfaces defined by expanded conformations. Nature Communications 13:7722.

95. Nag, A., G. Mitra, and P. C. Ghosh. 1996. A colorimetric assay for estimation of polyethylene glycol and polyethylene glycolated protein using ammonium ferrothiocyanate. Analytical Biochemistry 237:224–231.

96. Palit, S., L. He, W. A. Hamilton, A. Yethiraj, and A. Yethiraj. 2017. Combining Diffusion NMR and Small-Angle Neutron Scattering Enables Precise Measurements of Polymer Chain Compression in a Crowded Environment. Physical Review Letters 118:097801.

97. Bolhuis, P., and D. Frenkel. 1994. Numerical study of the phase diagram of a mixture of spherical and rodlike colloids. The Journal of Chemical Physics 101:9869–9875.

98. Minton, A. P. 2020. Simple Calculation of Phase Diagrams for Liquid–Liquid Phase Separation in Solutions of Two Macromolecular Solute Species. The Journal of Physical Chemistry B 124:2363–2370.

